# GAMES: A dynamic model development workflow for rigorous characterization of synthetic genetic systems

**DOI:** 10.1101/2021.10.20.465216

**Authors:** Kate E. Dray, Joseph J. Muldoon, Niall M. Mangan, Neda Bagheri, Joshua N. Leonard

## Abstract

Mathematical modeling is invaluable for advancing understanding and design of synthetic biological systems. However, the model development process is complicated and often unintuitive, requiring iteration on various computational tasks and comparisons with experimental data. Ad hoc model development can pose a barrier to reproduction and critical analysis of the development process itself, reducing potential impact and inhibiting further model development and collaboration. To help practitioners manage these challenges, we introduce **GAMES**: a workflow for **G**eneration and **A**nalysis of **M**odels for **E**xploring **S**ynthetic systems that includes both automated and human-in-the-loop processes. We systematically consider the process of developing dynamic models, including model formulation, parameter estimation, parameter identifiability, experimental design, model reduction, model refinement, and model selection. We demonstrate the workflow with a case study on a chemically responsive transcription factor. The generalizable workflow presented in this tutorial can enable biologists to more readily build and analyze models for various applications.

**Graphical Abstract:** 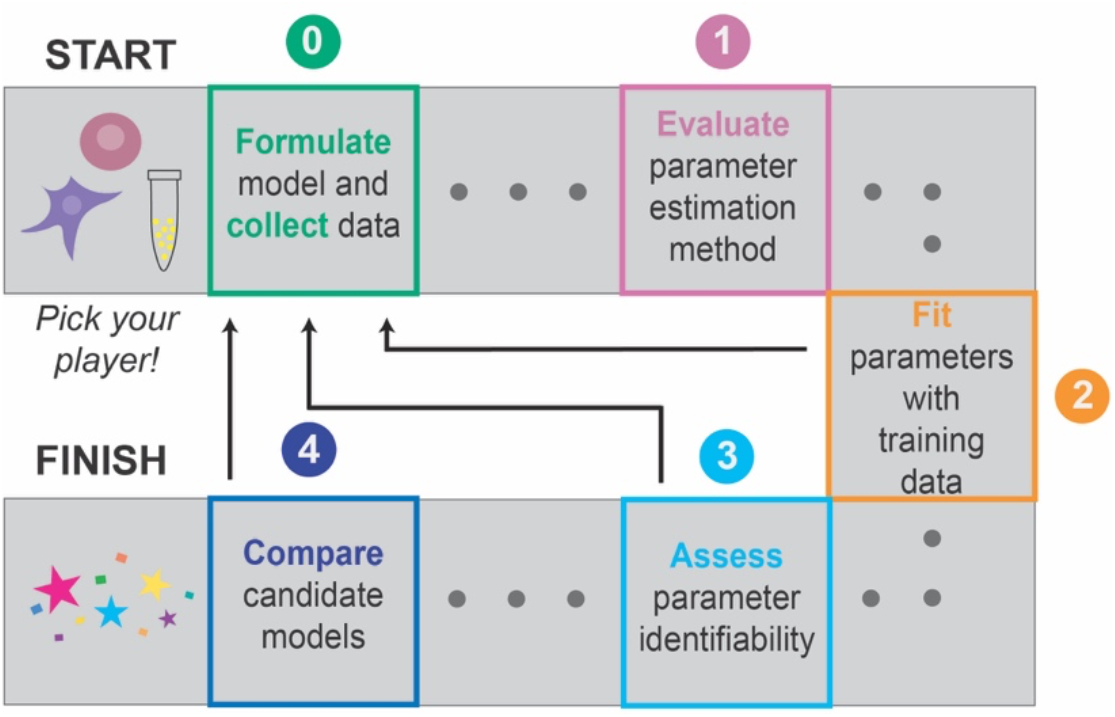

## Introduction

Dynamic mathematical models can be used to enable understanding, prediction, and control of biological systems. At a fundamental level, this type of model is a mathematical description of the components and interactions in a system. These models have proven useful for investigating the behaviors of genetic programs—sets of biological parts such as genes and proteins that are designed to implement specific functions^*1–5*^. Efforts to date have often used ordinary differential equations (ODEs), which describe the time evolution of the concentrations of system components and provide a framework for representing continuous dynamic systems. These models have been used to explain and predict behaviors of genetic programs that perform a growing array of functions including regulating gene expression^*3–7*^, implementing logic gates^*5, 6*^, and implementing feedback control^*8–13*^.

Models can generally be employed for *explanation*, in which the objective is to help the user understand a set of experimental observations, or for *prediction*, in which the objective is to simulate the response of a genetic program to a previously untested experimental condition or design choice (terms in italics are summarized in a glossary in **Supplementary Information**). When employed for explanation, models can help one understand how genetic programs work by identifying mechanisms that are necessary to describe a set of observations, or *training data*^*6, 11, 12*^. These models can help uncover mechanistic insight by testing whether a proposed model formulation is consistent with experimental observations, which may include data of different types and collected in separate experiments. Models employed for prediction build upon explanatory models (that are assumed to be accurate, at least locally) and aim to predict outcomes of a previously untested experimental condition or design choice, called *test data*. These models can guide genetic program design by predicting how relevant parameters and/or conditions can be adjusted to produce an intended response and by exploring how different well-characterized parts can be combined to carry out specific functions ^*3, 4, 7, 14, 15*^. Both usages can accelerate the “design, build, test, learn” cycle that is a paradigm in synthetic biology.

Although the value of model-guided design is generally recognized, the process of model development remains challenging, particularly for those without extensive training in investigating complex, multi-dimensional design spaces, dealing with unconstrained parameter spaces, and mapping experiments to simulations. The modeler must propose, implement, analyze, and refine candidate models until a suitable model is identified by making comparisons between model simulation outcomes and experimental observations. This iteration is often accomplished through intuition-guided exploration rather than through a reproducible and generalizable procedure. Ad hoc model development makes it challenging for both the modeler and any reader to evaluate how choices made during model development impact the utility of the model, and a lack of clarity makes it challenging to reproduce or extend this process to new applications. The lack of generally agreed-upon model development procedures might also contribute to the major challenge of limited reproducibility in biological models^*16, 17*^.

To help address these challenges, and to lower the barrier to entry for synthetic and systems biology researchers, we present a systematic workflow for model development. We focus our discussion on genetic programs, for which these approaches are well-developed, although the concepts extend to other settings. We demonstrate the workflow using a tutorial describing a simulation-based case study, which is a demonstration executed using simulated data that serve as a stand-in for experimental observations, and in which the model structure and parameters are known. This case study is based upon a hypothetical but realistic genetic program and a set of associated mechanistic assumptions, which are described in detail below. We include example code, which is written in the free and widely used programming language Python and available on GitHub, that executes each step of the workflow for the case study (**Methods, Supplementary Information**).

This tutorial is written with the assumption that the reader has familiarity with (1) identifying compelling biological questions to investigate through modeling and translating these questions into modeling objectives; (2) formulating ODE-based models that describe biomolecular systems based on mass action kinetics; and (3) collecting or otherwise obtaining supporting experimental data. For additional introduction to model formulation, we refer the reader to existing resources^*18, 19*^. Here we focus on topics that address the limitations of ad hoc model development, including methods for evaluating both the robustness of a parameter estimation method and the goodness and uniqueness of model fits to experimental data.

## Results

While there exists a deep body of knowledge that can guide the model development process, from model formulation^*18, 19*^ and parameter estimation^*19–25*^, to model selection^*23, 26, 27*^, parameter identifiability analysis^*25, 28–37*^, and experimental design^*7, 29, 31, 38, 39*^, navigating these tasks can be slow and cumbersome, posing a barrier to entry. This challenge is heightened by the fact that published studies often focus on a final optimized model rather than the process by which the model was generated, which is often through laborious iteration that is understandably difficult to fully capture in a report. A generalizable workflow to make model development more objective would improve rigor and reproducibility and lower the barrier to entry for synthetic and systems biology researchers. Towards this goal, we describe and demonstrate a rigorous process for developing and analyzing dynamic models, walking through methods and interrelated considerations for the steps of model formulation, parameter estimation, parameter identifiability analysis, experimental design, model reduction, model refinement, and model comparison and selection.

### A systematic, conceptual workflow for ODE modeling in synthetic biology

#### Workflow description

**GAMES**(**G**eneration and **A**nalysis of **M**odels for **E**xploring **S**ynthetic systems) describes the model development process as a set of five tasks, or modules (**Figure 1**). We introduce the overall logic of this workflow and then elaborate on key concepts and approaches within each module in subsequent sections. The modeler first uses Module 0 to initiate the process by defining the modeling objective, formulating one or more base case models, and collecting training data. In Module 1, a proposed *parameter estimation method* (PEM) is evaluated using simulated data from the model structure defined in Module 0. The PEM is then used in Module 2 to fit parameters to the training data. In this way, Module 1 ensures that the parameters are estimated using an appropriate method and computational implementation. If adequate agreement with the training data is obtained in Module 2, then parameter identifiability is assessed in Module 3. If all parameters are *identifiable* (capable of being uniquely estimated within a finite confidence interval), then candidate models can be compared in Module 4. Results from intermediate steps can motivate iteration (**Figure 1, dotted lines**) between experiments and simulations either to improve the agreement between experimental and simulated data or to constrain parameter estimates. The GAMES workflow helps the modeler track why and how parameters are tuned during model development and evaluate the extent to which this tuning is supported by data.

**Figure 1.**
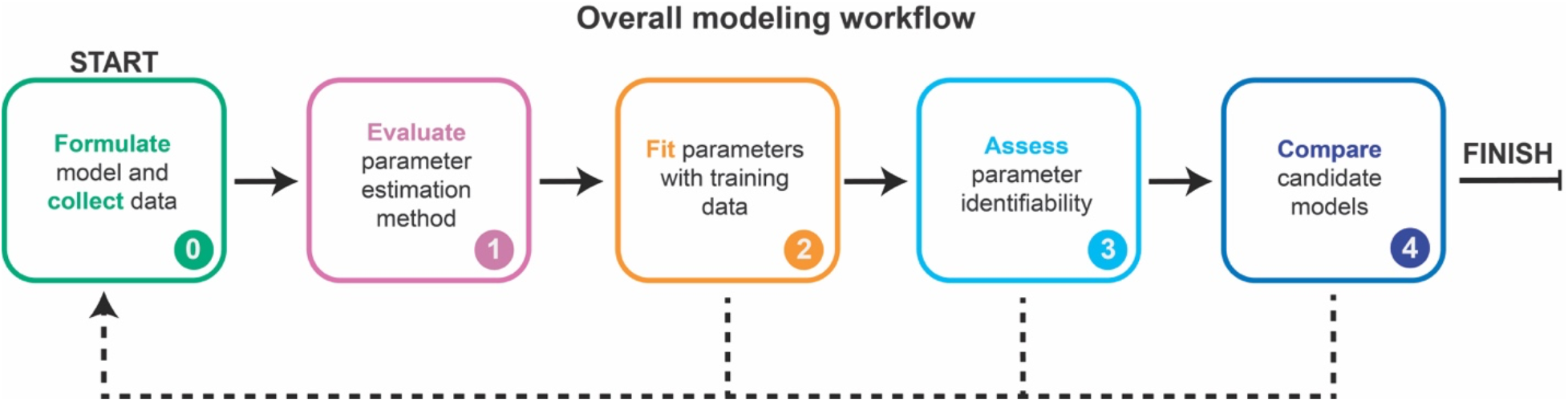
A systematic workflow for ODE modeling. Dotted lines represent iteration between modules, as further described in Figure 10.

#### Case study description

In the following sections, we demonstrate the GAMES workflow using a case study of a simple genetic circuit based upon a canonical synthetic biology part: a chemically responsive transcription factor (crTF) that regulates gene expression in mammalian cells. crTFs confer external control over the expression of a target gene in the presence of a chemical inducer. crTFs have been built using various mechanisms and genetic parts^*5, 6, 40, 41*^, and these systems can also confer titratable control over the expression of clinically relevant genes^*41*^. Typically, crTF protein-protein interactions are mediated by the association of cognate dimerization domains^*5, 6, 40, 41*^. Our hypothetical crTF has two protein parts, one containing a DNA-binding domain and the other containing an activation domain. In the presence of a ligand, these parts reconstitute at the interface of the dimerization domains. For simplicity, the term “DNA-binding domain” refers to the DNA-binding domain fused to one of the dimerization domains, and the term “activation domain” refers to the activation domain fused to the other dimerization domain. The reconstituted transcription factor can then activate a target promoter, leading to transcription and translation of a reporter protein that can be measured. We know that the doses of all three components affect the amount of reporter protein expressed with and without ligand, but the mechanism and parameters for the intermediate states are unknown and not measured. For the purpose of this case study, crTFs are a useful representative example of the types of states and interactions necessary to build a model of a genetic program, and the mechanism is simple but sufficiently nontrivial to demonstrate the utility of the workflow.

We first pose a *reference model* that reasonably describes a crTF system, so that we can run simulations of this reference model and generate data that serve as a stand-in for experimental observations. We refer to these reference model-generated data as training data in this tutorial to link this demonstration to the practical use of GAMES. We know the structure and parameters of the reference model, which includes reasonable mechanistic descriptions and physically plausible but essentially arbitrary *reference parameter* choices. In practice, one would have an experimental system rather than a reference model from which to collect data, and the model structure and parameters would be unknown. In these applications, the model structure would represent a set of mechanistic hypotheses about how the system works. To illustrate the workflow, we will use GAMES and the training data to perform parameter estimation and refinement on candidate models. We can then compare resulting parameter sets and simulated data to the known parameters of the reference model and the training data. We will first demonstrate the ideal workflow in which each module is successfully completed on the first try, then we will discuss common failure modes for each module and suggested actions for remediation (**Supplementary Table 1**) along with general paths for iteration between modules (**Figure 10**).

### Module 0: Collect data and formulate model(s)

#### Motivation and general method

The first challenge in model development is to identify the *modeling scope* (a choice of which aspects will be described or omitted based on the modeling objective(s)) and to use this choice to guide collection of base case training data and formulation of a base case model (**Figure 2a**, **Module 0.1**). Statistician George Box is credited with coining, “All models are wrong, but some are useful.” Defining the objective at the outset enables the modeler to identify what would make the model useful rather than trying to develop an unnecessarily complex model that may be difficult or even impossible to interpret.

**Figure 2.**
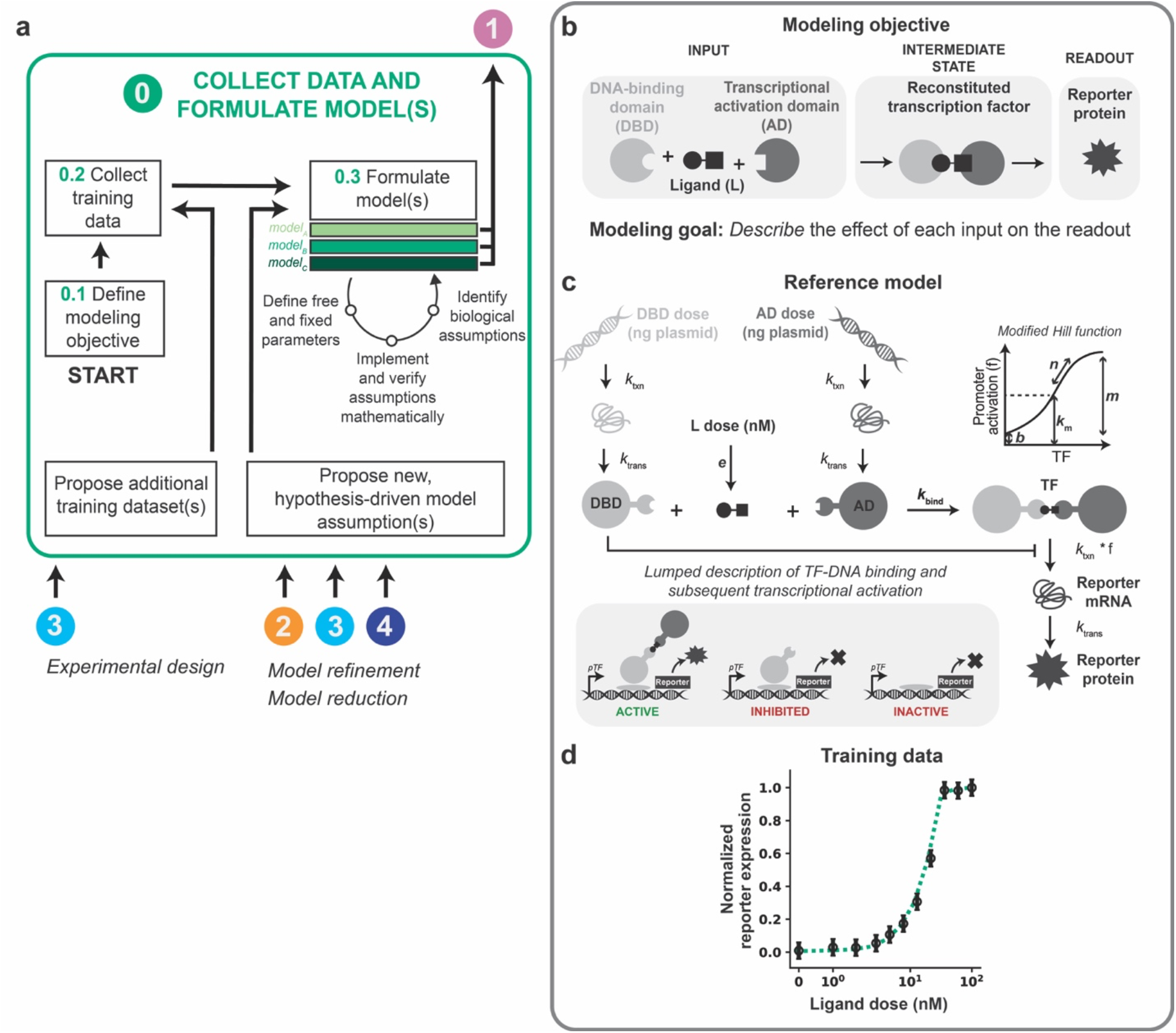
Collect data and formulate model(s). **(a–d)** This figure and subsequent figures present the overall workflow **(a)** and then examine its application to the case study **(b–d)**. **(a)** Module 0 workflow for collecting training data and formulating model(s). **(b–d)** Module 0 case study for a hypothetical crTF. **(b)** Case study modeling objective. The crTF has a DNA-binding domain and an activation domain that reconstitute in the presence of a ligand. The DNA-binding domain and activation domain are each fused to a heterodimerization domain. The reconstituted crTF induces transcription of a reporter gene, and the mRNA translated into a protein that is measured. **(c)** Schematic of the reference model for the case study. Free parameters are in bold and other parameters are fixed based on literature values (**Supplementary Information**). Promoter activation is described by a modified Hill function that accounts for these promoter states: active (bound by crTF), inhibited (bound by DNA-binding domain), and inactive (unbound). Degradation reactions are included for all states and for simplicity are not depicted here. Reference parameter values are in **Supplementary Table 2**. **(d)** Case study training data. The initial set of training data is a ligand dose response at a constant dose of DNA-binding domain plasmid and activation domain plasmid (50 ng of each plasmid per well of cells).

The modeling objective will guide decisions on the collection of training data (**Figure 2a**, **Module 0.2**) and formulation of one or more base case models (**Figure 2a**, **Module 0.3**). Together, the modeling objective, training data, and base case model define the modeling problem. Models differ in biological assumptions, mathematical implementation of assumptions, and/or the definition of free and fixed parameter values or bounds. In this way, different models can represent different mechanistic hypotheses^*19*^. If there exists prior knowledge on physically realistic bounds for any parameters, this information should be included prior to parameter estimation. For example, if a model includes a first-order rate constant describing the rate of protein degradation as a free parameter, one could use knowledge from literature (depending on the organism and protein) to determine realistic bounds for this parameter.

Training data and reference model data are normalized to their respective maximum values. Data points and error bars are the simulated training data with added noise. The dotted line is the reference model. Data are plotted on symmetrical logarithmic (“symlog”) axes in this figure and all subsequent figures for ligand dose response simulations.

#### Case study

We define the modeling objective to describe the effect of each input (ligand dose, DNA-binding domain plasmid dose, AD plasmid dose) on the readout (reporter protein expression) (**Figure 2b**). In other scenarios, the objective might be both to explain a set of observations (training data) and to predict another set of observations (test data). We limit the scope of this example to an explanatory objective.

The reference model (**Equations 1–8, Figure 2c**) describes transcription, translation, and binding of the inputs, activation of the reporter promoter, and transcription and translation of the reporter. Constitutive transcription and translation of the DNA-binding domain and the activation domain are described by mass action kinetics with rate constants k_txn_ for transcription and k_trans_ for translation. Basal degradation of the DNA-binding domain and activation domain are described by first-order rate constants k_deg,m_ for mRNA species and k_deg,P_ for protein species. Transcription, translation, and degradation rate constants for states of the same type (e.g., mRNA or protein) are constant and equal except for the degradation rate constant of the reporter protein (**Supplementary Table 1**).

Abbreviations — DBD: DNA-binding domain, AD: activation domain, L: ligand, m: mRNA, p: protein, txn: transcription, trans: translation

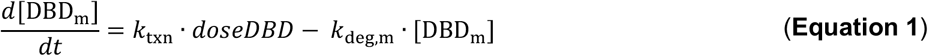

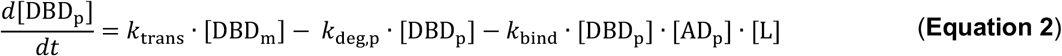

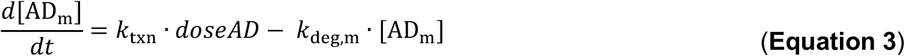

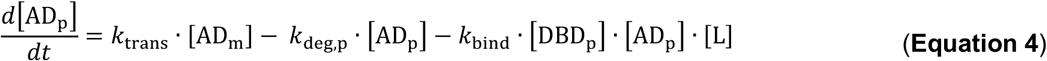

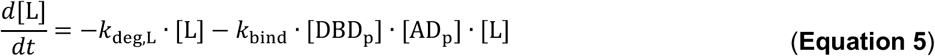

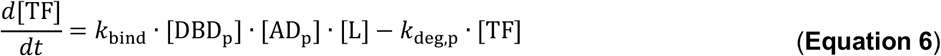

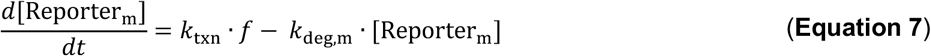

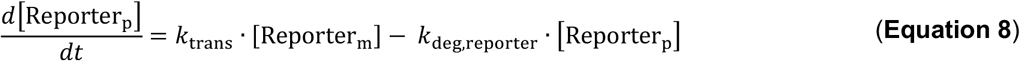

The ligand dose is multiplied by a parameter *e* (a lumped parameter representing a conversion factor between the ligand dose in experimental units and arbitrary units used in the model and also the effective diffusion rate of the ligand into a cell) and then used as the initial condition for the ligand state, as described by **Equation 9**.

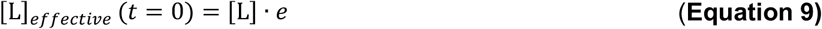

Ligand degradation is described by a first-order reaction with a rate constant k_deg,L_. This formulation includes the assumption that the three inputs (DNA-binding domain protein, activation domain protein, and ligand) bind in an irreversible, trimolecular reaction (a choice that simplifies the model for the purposes of this tutorial) described by the rate constant k_bind_.

The reconstituted transcription factor, TF, can then induce promoter activation. The DNA-binding domain is a competitive inhibitor of TF binding to DNA, as the DNA-binding domain and TF each bind to the same promoter region. A modified Hill function that is commonly used to represent a transcriptional activator and a transcriptional inhibitor acting at the same promoter^*42*^ (**Equation 10**) was used to lump together transcription factor and DNA (TF-DNA) binding (with the possibility of cooperativity in binding) and subsequent transcription of the reporter. This function, *f*, takes into account the promoter states: active (bound by TF), inhibited (bound by DNA-binding domain), and inactive (unbound).

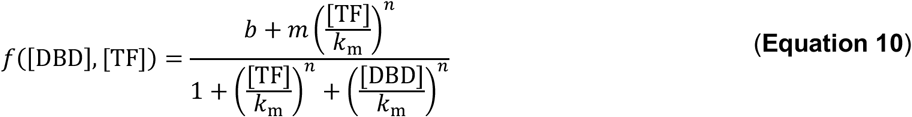

Background promoter activation, without activator or inhibitor present, is described by *b*. Maximal expression from the promoter, at a saturating amount of TF, is described by *m*. The activation coefficient *k_m_* for the crTF is assumed equal to the inhibition coefficient for the inhibitor. The Hill cooperativity *n* applies to cooperative TF-DNA binding. Reporter mRNA is generated according to the promoter activation function described above multiplied by a first-order rate constant k_txn_, and degradation is described by a first-order rate constant k_deg,m_. Reporter protein is generated based on a first-order rate constant k_trans_, and degradation is described by a first-order rate constant, k_deg,reporter_.

There are six free parameters—*e*, *k*_bind_, *b*, *m*, *k_m_*, and *n*— and the other parameters are fixed based on literature values (**Supplementary Table 1**). Reference parameter values were chosen arbitrarily from within a plausible range (**Supplementary Table 2**). The bounds for each free parameter except *n* were set to three orders of magnitude in either direction of the reference value. The bounds for *n* (the Hill coefficient for cooperative TF-DNA binding) are 1 and 4, which are considered relevant values^*42*^. A useful resource for choosing parameter estimates is BioNumbers^*43*^, a database of salient biological numbers curated from the literature.

To generate a set of training data akin to experimental data, we considered a scenario in which ligand dose (independent variable) was varied between 0–100 nM (11 datapoints including 0, and 10 logarithmically spaced datapoints between 1–100 nM) while holding plasmids encoding DNA-binding domain and activation domain constant at 50 ng each per well of transfected cells. The dependent variable is the reporter protein at 24 h after ligand treatment (**Methods**). Training data were generated by simulating the reference model with the reference parameters for these conditions (**Figure 2d**). Each datapoint was normalized to the maximum value within the data set (**Methods**).

In constructing the training data set, we separately incorporated the effects of technical error and biological variation. To represent biological variation, error bars were added to each data point. The error bars represent a distribution of values one might observe if the experiment were repeated many times with different biological replicates. Error bars on all plots represent the standard deviation, which was set to a constant arbitrary value for each data point. To represent technical error, randomly generated noise was added to each data point based on normally distributed error described by *N*(0, *σ_SEM_*^2^), where *σ_SEM_* is the standard error of the mean. This error is the inaccuracy associated with the measurement of each data point in an analogous experiment. Representing error and variation in this way approximates phenomena in experimental training data. Now that we have a modeling objective, a training data set, and a base case model (that matches the reference model in formulation for the purpose of the case study but is not yet parameterized), we are ready to apply the GAMES workflow.

### Module 1: Evaluate parameter estimation method

#### Motivation

Given a base case model and training data, we next aim to identify the set of parameters that produce the best agreement between this model and data set. This objective can be accomplished using a parameter estimation method (PEM): an algorithm for identifying the set (or sets) of parameters that yield the best agreement between training data and simulations of a model of interest. Maximizing agreement is analogous to minimizing a predefined *cost function* that accounts for the difference between each training data point and the corresponding simulated value. The parameter set yielding the lowest cost is the *calibrated parameter set*. In addition to defining the cost function, several PEM covariates and hyperparameters must be appropriately defined. A *covariate* is an aspect of the model development process, which results from choices made by the modeler, and that is not of primary interest but can impact the interpretation of results. Covariates include the choices of scale and constraints of parameters, ODE solver, optimization algorithm, and user-friendliness of the code^*20*^. Algorithm-specific *hyperparameters* must be specified to set up, execute, and interpret a PEM^*20*^. For example, for a global search (a high-dimensional sweep over defined regions of parameter space), the number of parameter sets is a hyperparameter, and for a multistart optimization algorithm, the number of initial guesses is a hyperparameter.

It is important to select a PEM that is well-suited to the *parameter estimation problem*, which is defined by the training data, model, and cost function (**Figure 3a**). The cost function landscape, which represents the relationship between parameter values and the cost function, is unique and unknown a priori. The PEM explores this landscape to find an optimal parameter set or sets. The simplest cost function landscapes in 1D parameter space have a single local minimum. However, as either the number of local minima or number of dimensions in parameter space increase, the cost function landscape becomes more difficult to explore and the global minimum (or minima) is harder to find. Cost function landscapes with flat regions, in which the model is unresponsive to changes in parameters, are also difficult to explore, as the algorithm can get lost wandering around these regions and fail to converge on the global minimum^*44*^. Flat cost functions are common in biological systems modeling because systems are often only partially accessible, meaning that only a few states are observed and used as training data, and therefore timescales of intermediate reactions are often not well-constrained^*31*^.

**Figure 3.**
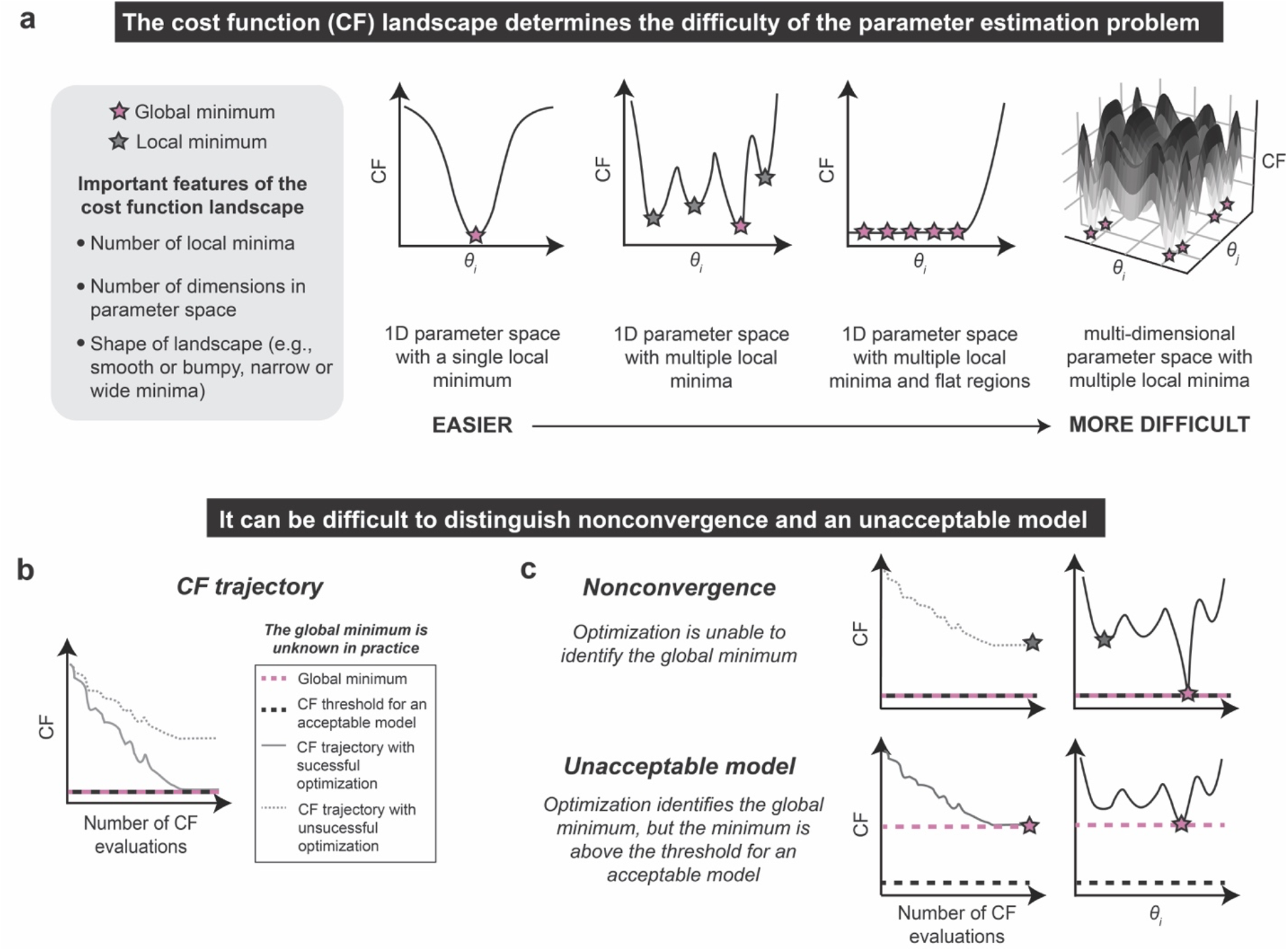
The parameter estimation method should be well-suited to the parameter estimation problem. **(a)** Relationship between cost function shape and difficulty of parameter estimation. *θ_i_* is a free parameter. In the rightmost plot, the shading corresponds to the value of the cost function (z-axis). **(b, c)** Distinction of nonconvergence and an unacceptable model. **(b)** A cost function trajectory indicates the value of the cost function at each iteration of the optimization algorithm. A successful optimization will identify a global minimum, although it will not necessarily identify all global minima. An acceptable model (i.e., parameter set) will yield a cost function value that is less than or equal to some threshold. **(c)** If the optimization algorithm does not converge (top), then global minima of the model (pink) cannot be found and instead a local minimum (gray) is identified. If the model is unacceptable (bottom), then a global minimum of the cost function is found (pink), but this minimum is too large to meet the cost function threshold predetermined to define an acceptable model (black dotted line). In practice, the global minimum is unknown and therefore these situations are indistinguishable (or could occur simultaneously).

When analyzing parameter estimation results, it can be difficult to distinguish the scenarios of nonconvergence and having an unacceptable model. If the optimization algorithm is successful, then the cost function trajectory will converge to a global minimum (**Figure 3b**). If that global minimum is below the threshold set to define an acceptable model, then the model is considered sufficient. However, if the optimization converges to a local minimum with a cost above the defined threshold, then the optimization is not successful. The global minimum is not known a priori, and therefore it is often not possible to distinguish the scenarios of nonconvergence and having an unacceptable model (**Figure 3c**). This challenge arises because the cost function trajectories for each of these scenarios may look nearly identical, and it is only with knowledge of the global minimum (knowledge that we do not have in practice) that the two cases can be distinguished. Therefore, it is wise to perform consistency checks where the global minimum is known to assess whether a candidate PEM is appropriate for the given problem and is implemented properly, enabling identification a global minimum of the model-specific cost function. The consistency check serves as a positive control for parameter estimation, paralleling common positive controls used in wet lab experiments to ensure that an assay was functional and was correctly executed. Below, we consider how a consistency check can be performed and interpreted for a proposed PEM.

#### General method

##### Parameter estimation method

In the absence of knowledge about features of the cost function landscape, a reasonable choice of PEM is multi-start optimization using the Levenberg-Marquardt optimization algorithm^*45*^ (**Figure S1**). We implemented this method as follows: 1) execute a Latin hypercube global search over parameter space^*46*^ to generate n_search_ parameter sets; 2) filter the results by cost function and choose the top n_init_ parameter sets; 3) run n_init_ optimization simulations using the Levenberg-Marquardt algorithm^*47, 48*^, each time using a different parameter set as an initial guess. The parameter set with the lowest cost function after optimization is considered the best parameter set. We use the error-weighted *χ^2^* as the cost function (**Equation 11**).

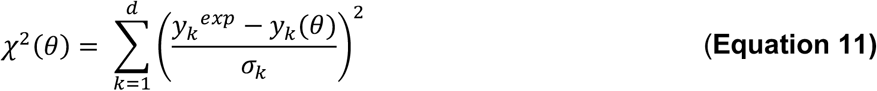

Here, *d* is the total number of datapoints in the training data set, y_k_^exp^is the k^th^ datapoint in the experimental data with associated measurement error (standard deviation) *σ_k_*, and *y_k_*(*θ*) is the simulated value of the k^th^ datapoint using the parameter set *θ*. This cost function is appropriate for error that is approximately normally distributed; if the standard deviation is not normally distributed, then a different metric such as the root mean squared error can be used instead. For this PEM, covariates include the global search and optimization algorithms and the choice of cost function, and hyperparameters include n_search_ and n_init_. A different PEM might be required depending on the model structure, training data, and cost function. For example, if multiple objectives are specified, such as for fitting data describing more than one species or for multiple independent experiments of different types, and as contrasted with a single objective in Equation 11, then one option is to use an evolutionary algorithm to conduct multi-objective optimization^*49*^.

##### Consistency check

We can test the proposed PEM by performing a consistency check to assess whether the PEM finds the global minima when the parameters are known^*20*^. We use the model structure defined in Module 0 and choose a set of parameter values to generate a PEM evaluation data set. PEM evaluation data are generated for the specific and limited purpose of assessing whether the PEM and hyperparameters selected are suitable in general for fitting models that have the same structure as the model defined in Module 0. The PEM evaluation parameters should be chosen such that PEM evaluation data sets qualitatively match the training data, if possible. Finding PEM evaluation parameters can be accomplished by performing a global search over the free parameter space, calculating the cost function with respect to the training data for each parameter set, and choosing the parameter sets with the lowest cost functions as PEM evaluation parameters. Synthetic noise similar to the expected measurement error in the training data should also be applied, and data should be re-normalized, if applicable, afterwards. We then evaluate the performance of the PEM based on whether the method can identify parameter sets yielding acceptable agreement between the simulated data and each PEM evaluation data set (**Figure 4a, Module 1.1**). PEM evaluation problems are useful because we know that in each case, an acceptable solution (a high-performing parameter set) does exist. Therefore, if the PEM cannot identify an acceptable solution for a PEM evaluation problem, then we assume that the method will also be unable to identify an acceptable solution when applied to the training data. It is good practice to include multiple PEM evaluation data sets to represent a variety of plausible model time courses and local minima, each of which might pose unique challenges for parameter estimation.

**Figure 4.**
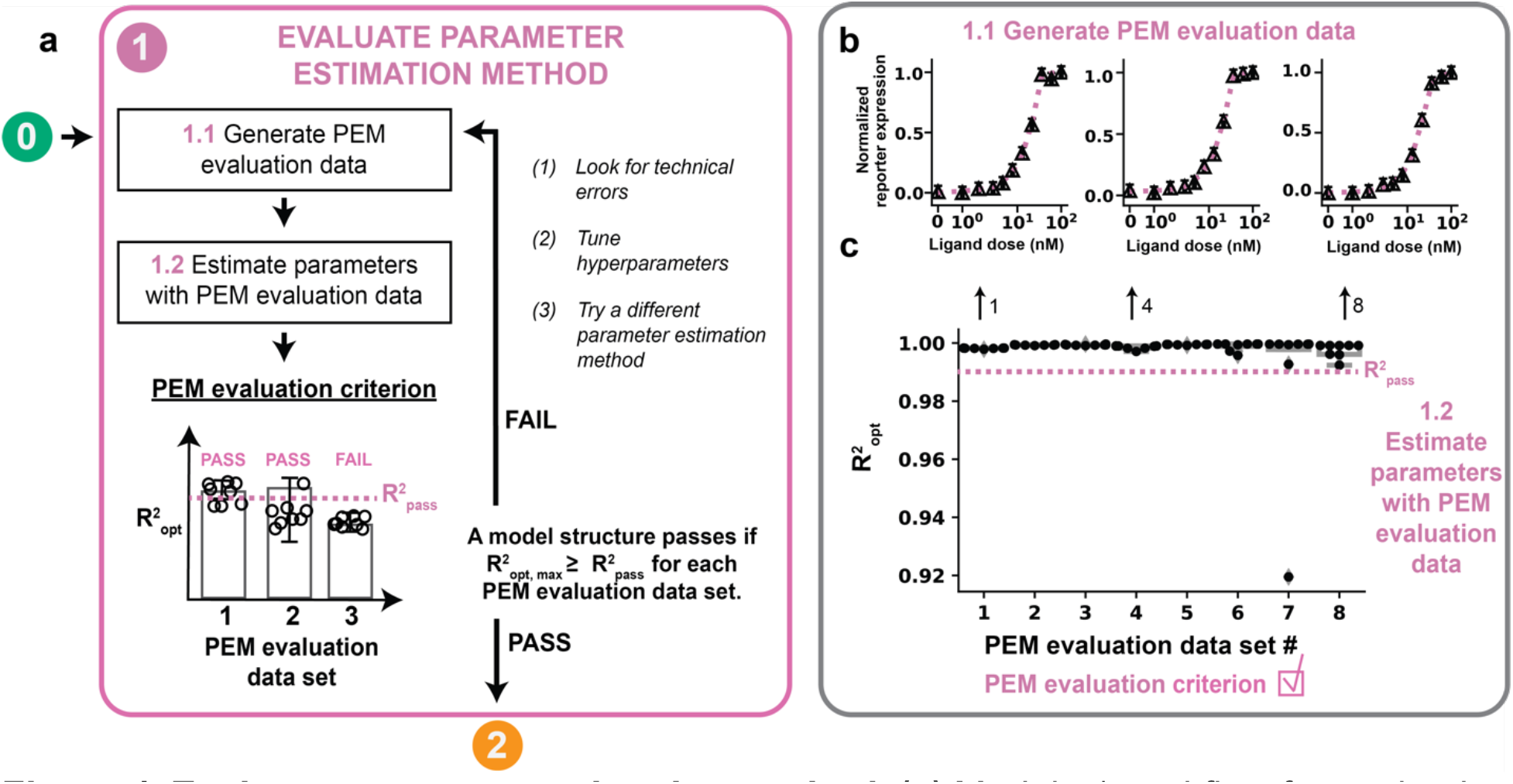
Evaluate parameter estimation method. **(a)** Module 1 workflow for evaluating the PEM using simulated training data. A model must pass the PEM evaluation criterion before moving on to Module 2. **(b, c)** Module 1 case study for a hypothetical crTF. **(b)** Generating the PEM data. A global search of 1000 parameter sets was filtered by *χ*^2^ with respect to the training data and the 8 parameter sets with the lowest *χ^2^* values were used as reference parameters to generate PEM evaluation data. For each data set, technical error was added using a noise distribution of *N*(0, 0.017^2^). Triangle datapoints are PEM evaluation data. **(c)** Determination of the PEM evaluation criterion. For each PEM evaluation data set, a global search with 100 randomly chosen parameter sets was used to choose 10 parameter sets to use as initial guesses for optimization. The optimized parameter sets and cost function from each of the PEM evaluation problems were used to evaluate the PEM evaluation criterion. Each parameter was allowed to vary across three orders of magnitude in either direction of the reference parameter value, except for *n*, which was allowed to vary across [10^0^, 10^0.6^]. Results are shown only for parameter sets yielding *χ^2^* values within the bottom (best) 10% of *χ^2^* values (to the left of the pink dotted line in **Figure S2b**) achieved in the initial global search with respect to the training data (Module 1.1). Only parameter sets yielding R^2^ > 0.90 are included on the plot to more clearly show data points with R^2^ values that exceed R^2^_opt_. Both of these filtering strategies apply to all plots of PEM evaluation data in this tutorial.

The consistency check is evaluated by a quantitative criterion, such as the coefficient of determination, R^2^, to measure the ability of the method to solve the PEM evaluation problems (**Equation 12**).

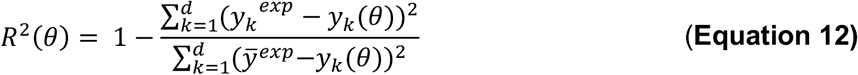

In the formula, *d* is the number of data points in the training data set, *y_k_^exp^* is the k^th^ data point in the experimental data with associated measurement error (standard deviation) *σ*_k_, *y*_k_(*θ*) is the simulated value of the k^th^ data point using the parameter set *θ*, and *ȳ^exp^* is the mean of the training data set. R^2^ is used to evaluate the correlation between normalized training data values and normalized model-simulated values (**Figure S2a**). If the criterion meets some pre-determined threshold, then the method is considered appropriate. Here, the PEM evaluation criterion is satisfied if any parameter set yielding an R^2^ of at least 0.99 (defined as R^2^_pass_) can be identified for each PEM evaluation problem (**Figure 4a, Module 1.2**). The criteria should be stringent enough to determine if the known PEM evaluation parameters were found within the known noise that was added to make the PEM evaluation data similar to the training data. For example, the R^2^ between each PEM evaluation data set with and without noise can be calculated, and R^2^_pass_ can be set to the mean of these values. If the PEM evaluation criterion is satisfied, then the modeler can move on to Module 2. If the criterion is not satisfied, then the modeler should look for technical errors such as in the implementation of the PEM, tune the hyperparameters and/or covariates, or try a different PEM until the criterion is satisfied. For example, if R^2^ < R^2^_pass_, then it is possible that a local minimum has been identified, which suggests that different hyperparameters or covariates should be used to reach a global minimum. The PEM evaluation criterion should be determined for each new version of the model, because changes in model structure (i.e., formulation) or choice of training data could impact the ability of the PEM to identify an acceptable solution.

Quantifying goodness of fit with R^2^ has some limitations, as the metric quantifies the correlation between training data and simulated values, not direct agreement between the experimental and simulated data, so the correlation between R^2^ and *χ*^2^ should be validated by simulation studies and by visual inspection. One limitation is that R^2^ can be used to quantify the PEM evaluation criterion only in regimes in which R^2^ and *χ*^2^ are correlated, which can be validated by plotting R^2^ and *χ*^2^ values for simulations using randomly generated parameter sets (**Figure S2b**). Observing such a correlation enables the modeler to use R^2^ as a goodness of fit metric. If the correlation is not observed, the modeler would not be able to assume that parameter sets yielding high R^2^ values would also yield low *χ*^2^ values and could therefore not use R^2^ as a goodness of fit metric. The modeler should also visually inspect the best model fits to the PEM evaluation data to check whether parameter sets yielding high R^2^ values produce simulations that match the training data. Although it is possible to use *χ^2^* to define the consistency check, in practice one would need to adjust the value of *χ^2^_pass_* on a case-by-case basis, as the range of possible *χ^2^* values is dependent on the number of data points, the normalization strategy, and the measurement error. If R^2^ and *χ^2^* are not correlated, the PEM evaluation criterion could be defined by *χ*^2^, which could be defined by evaluating the *χ*^2^ values associated with agreement between the PEM evaluation data with and without simulated noise, as described for R^2^ in the previous paragraph.

#### Case study

Module 1 was executed, and the PEM evaluation criterion was satisfied. A global search with 1000 randomly chosen parameter sets was filtered by *χ*^2^ with respect to the reference training data, and the 8 parameter sets with the lowest *χ*^2^ values were used to generate 8 unique sets of PEM evaluation data (**Figure 4b**, **Figure S3**). To ensure the PEM evaluation data are as similar as possible in structure to the training data, biological and technical variation were added to each PEM evaluation data point using the method described in Module 0. The hyperparameters n_search_ and n_opt_ were set to 100 and 10, respectively. For each PEM evaluation problem, the resulting 10 optimized parameter sets were used to evaluate the PEM evaluation criterion. R^2^_pass_ was determined by calculating the mean R^2^ between each PEM evaluation data set with and without noise (R^2^_pass_ = 0.999, rounded down to R^2^_pass_ = 0.99). At least one parameter set yielding an R^2^ > R^2^_pass_ was identified for each PEM evaluation problem so the PEM evaluation criterion was satisfied (**Figure 4c**).

Our simulation study shows that R^2^ and *χ*^2^ are correlated only for relatively small values of *χ*^2^ (**Figure S2b**). At high values of *χ*^2^, high R^2^ values are sometimes still attainable when simulations are flat lines (no change in reporter expression with changes in ligand dose). This is because the R^2^ evaluates the difference between how well the simulated data from the model describe the experimental data (**Equation 12**, numerator) and how well the mean of the experimental data describes the experimental data (**Equation 12**, denominator). To ensure that R^2^ is evaluated only in a regime in which R^2^ and *χ*^2^ are correlated, we evaluate R^2^ only for parameter sets yielding *χ*^2^ values in the bottom (best) 10% of *χ*^2^ values (pink dotted line in **Figure S2b**) achieved in the initial global search with the training data. Visual inspection of the model fits to the PEM evaluation data (**Figure 4b**) shows that each data set is appropriately described.

### Module 2: Fit parameters with training data

#### Motivation

We can now use the selected and evaluated PEM to fit parameters to the training data and determine whether the resulting model adequately matches the training data. Based on the consistency check in the previous module, we have confidence that this approach will yield a parameter set corresponding to a global minimum of the cost function. If the model is acceptable, this global minimum will correspond to a good fit to the training data. If the model is not acceptable, this global minimum will not correspond to a good fit to the training data.

#### General method

To evaluate the best fit (or fits if multiple optimized parameter sets yield similar *χ*^2^ values), the training data and corresponding simulations are visually inspected to assess whether the model sufficiently fits each relevant feature of the training data (**Figure 5a**). The modeling objective determines which features must be captured in a sufficient fit (and, if applicable, which features need not be captured). For example, if the goal is to qualitatively recapitulate a set of observations, and the model is not expected to yield near-perfect agreement with the training data (perhaps because the modeler is not attempting to represent a very granular mechanism), then perhaps some quantitative features of the data can be ignored. At this stage, the dynamic state trajectories should also be plotted and inspected for physical plausibility. For example, if any trajectories have negative values, there is likely a numerical instability or implementation error that should be addressed before proceeding.

**Figure 5.**
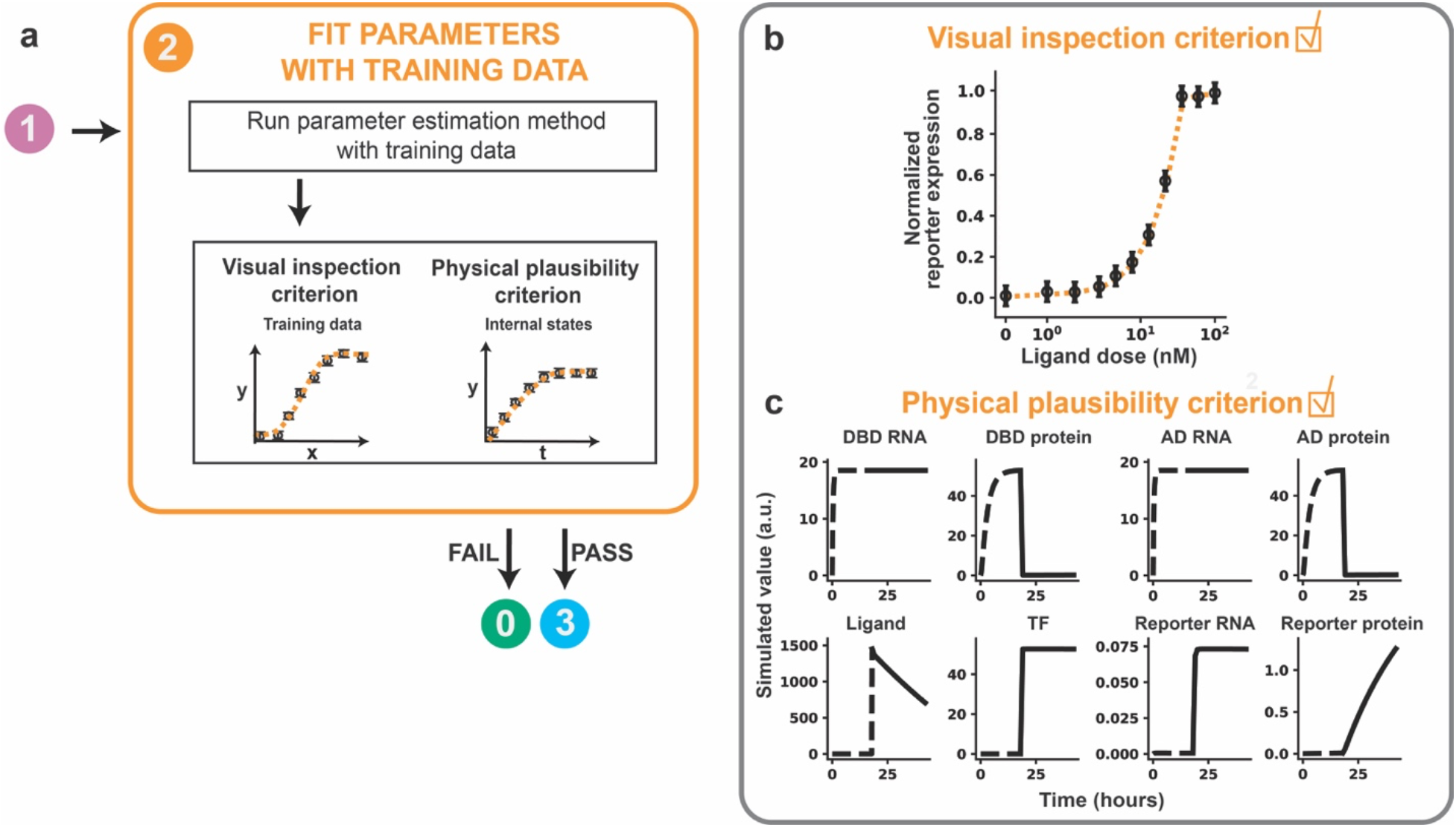
Fit parameters with training data. **(a)** Module 2 workflow for fitting parameters to training data. A model must pass both criteria before the modeler proceeds to Module 3. **(b, c)** Module 2 case study for a hypothetical crTF. **(b)** Visual inspection criterion. A global search with 100 randomly chosen parameter sets was used to choose 10 initial guesses for optimization. Each parameter was allowed to vary across three orders of magnitude in either direction of the reference parameter value, except for *n*, which was allowed to vary across [10^0^, 10^0.6^]. The parameter set with the lowest value of the cost function was chosen as the calibrated parameter set and is shown. Calibrated values are in **Supplementary Table 2**. **(c)** Physical plausibility criterion. Time course trajectories of each state variable in the reference model are shown for the highest ligand dose (100 nM). Dotted lines represent time before ligand treatment, and solid lines represent time after ligand treatment. Each state variable is in distinct arbitrary units, and thus values should not be compared between state variables. However, for any state variable, the trajectories can be compared across simulations, e.g., with different parameter values.

If the model fails to meet either the visual inspection criterion or physical plausibility criterion, the model is not a reasonable candidate. If the model fails the visual inspection criterion, the modeler can propose an updated model structure to reflect a new mechanistic hypothesis and return to Module 0 with the new model. If the model fails the physical plausibility criterion, the modeler can check the model implementation and consider options for resolving numerical instability, such as rescaling equations, trying a stiff ODE solver, or changing integration error tolerances ^*20, 50*^. This cycle is repeated, and once both criteria are satisfied, the modeler can proceed to Module 3.

#### Case study

Module 2 was executed for the case study, and the fitting criteria were satisfied. The parameter set with the lowest *χ*^2^ yielded an R^2^ value of 0.999. Inspection of the simulated training data showed that the model passed the visual inspection criterion because all features of the data, including the background reporter expression (without ligand treatment), steepness of the response curve, and saturating ligand dose are well-described (**Figure 5b**). The internal model state dynamics at the highest ligand dose (100 nM) satisfy the physical plausibility criterion, as each trajectory is physically realistic and does not include numerical instabilities or negative state variables (**Figure 5c**).

### Module 3: Assess parameter identifiability

#### Motivation

The modeler now has an estimate of parameter values that yield good agreement between the training data and the model. However, these values might not be unique, as other parameter sets might produce similar agreement. This ambiguity is a recurring challenge in ODE model development because if the parameters are not well-constrained, then it is also possible that downstream model predictions and conclusions will not be well-constrained. This issue arises when the training data do not fully constrain the parameters, such as if there are timescales that are too fast to measure, intermediate states are not observed, or the model is too complex given the available data. Unconstrained parameters often cause problems when making predictions because changes in a free parameter *θ_i_* might not affect the fit of the model to the training data, but the changes could affect predictions (i.e., simulated test data). For example, 8 of the 10 optimized parameter sets from the previous module (Module 1 case study) yield R^2^ > 0.99 and, and by visual inspection each of these optimized parameter sets fit the training data similarly well (**Figure 6a**); however, plotting the distribution of optimized values for each parameter shows that the values for most of parameters are not unique (**Figure 6b**). Therefore, many different parameter values yield similar agreement with the training data This result suggests that most of the parameters are unidentifiable—a phenomenon associated with flat cost function landscapes in which changes in a parameter *θ_i_* do not affect the value of the cost function in some or all regions of parameter space (**Figure 6c**). Ideally, the model will be at an appropriate level of complexity to describe the data, resulting in a model without unidentifiable parameters. In this module, we first focus on evaluating parameter identifiability before considering how to handle unidentifiable parameters should a model not exhibit the aforementioned ideal behavior.

**Figure 6.**
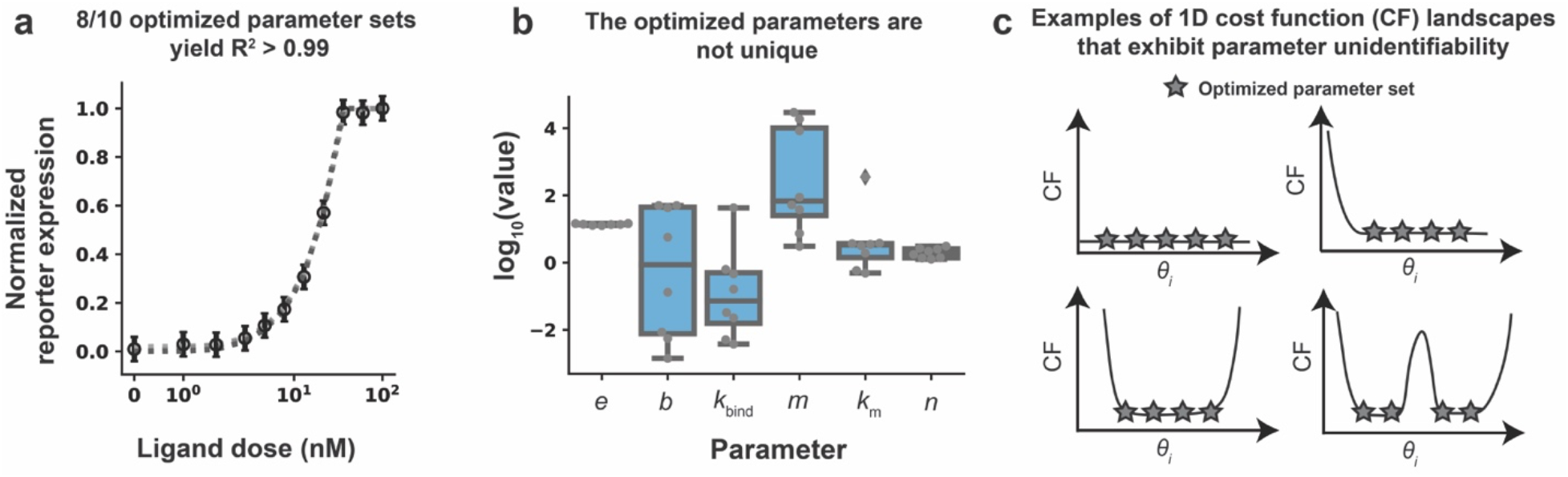
Many unique parameter sets describe the training data equally well. **(a)** Comparison across 8 optimized parameter sets identified in Module 1 (case study) that each yield R^2^ > 0.99. Simulated data generated with each of these parameter sets are shown in shades of gray (simulations overlap). **(b)** Distribution of each optimized parameter value. Each data point is an optimized parameter value. Most parameter values vary widely across multiple orders of magnitude while retaining very similar R^2^ values. **(c)** Examples of cost function landscapes exhibiting parameter unidentifiability. A parameter is considered unidentifiable if it cannot be uniquely estimated.

#### General method

A systematic way to determine whether parameter values are well-constrained given the training data is the parameter profile likelihood (PPL), which classifies parameters as identifiable, practically unidentifiable, or structurally unidentifiable^*30, 31, 34, 38, 51*^ (**Box 1**). Specifically, the PPL determines whether each parameter is appropriately constrained given the training data and measurement error. The PPL is determined individually for each parameter *θ_i_* by fixing *θ_i_*, re-optimizing all other free parameters *θ_j≠i_*, and calculating the minimum achievable *χ*^2^ (**Box 1 Figure B1a**). In this way, *χ*^2^ is kept as small as possible along *θ_i_*. This process is repeated for different values of *θ_i_* (**Box 1 Figure B1a**) until a confidence threshold *χ_PL_*^2^ = Δ_1-*α*_ is reached (**Box 1 Figure B1b**). The confidence threshold enables determination of the range of parameter values supported by the training data by accounting for measurement error associated with the training data. Parameter sets yielding PPL values below Δ_1-*α*_ are considered not significantly different from one another in terms of agreement with the training data. The shape of each individual PPL is used to classify each parameter as identifiable, structurally unidentifiable, or practically unidentifiable (**Box 1 Figure B1c**). Identifiable parameters reach Δ_1-*α*_ in both the positive and negative directions, practically unidentifiable parameters do not reach Δ_1-*α*_ in at least one direction, and structurally unidentifiable parameters have flat PPLs. Following classification of each parameter, one can explore the relationship between each unidentifiable parameter and other parameters, and the relationship between each unidentifiable parameter and internal model states, predictions, or other relevant observables defined in the modeling objective (**Box 1 Figure B1d**). We use the term “model predictions” from this point forward for relevant observations made using the model, regardless of whether test data are considered. After each parameter is classified, the PPL approach includes paths for model reduction and experimental design^*30, 37, 34, 38, 57*^ to propose and evaluate refinements to the model to improve identifiability. The steps for assessing parameter identifiability using the PPL are summarized in **Figure 7a**.

##### Box 1: Parameter identifiability analysis via evaluation of the parameter profile likelihood

**Figure B1.**
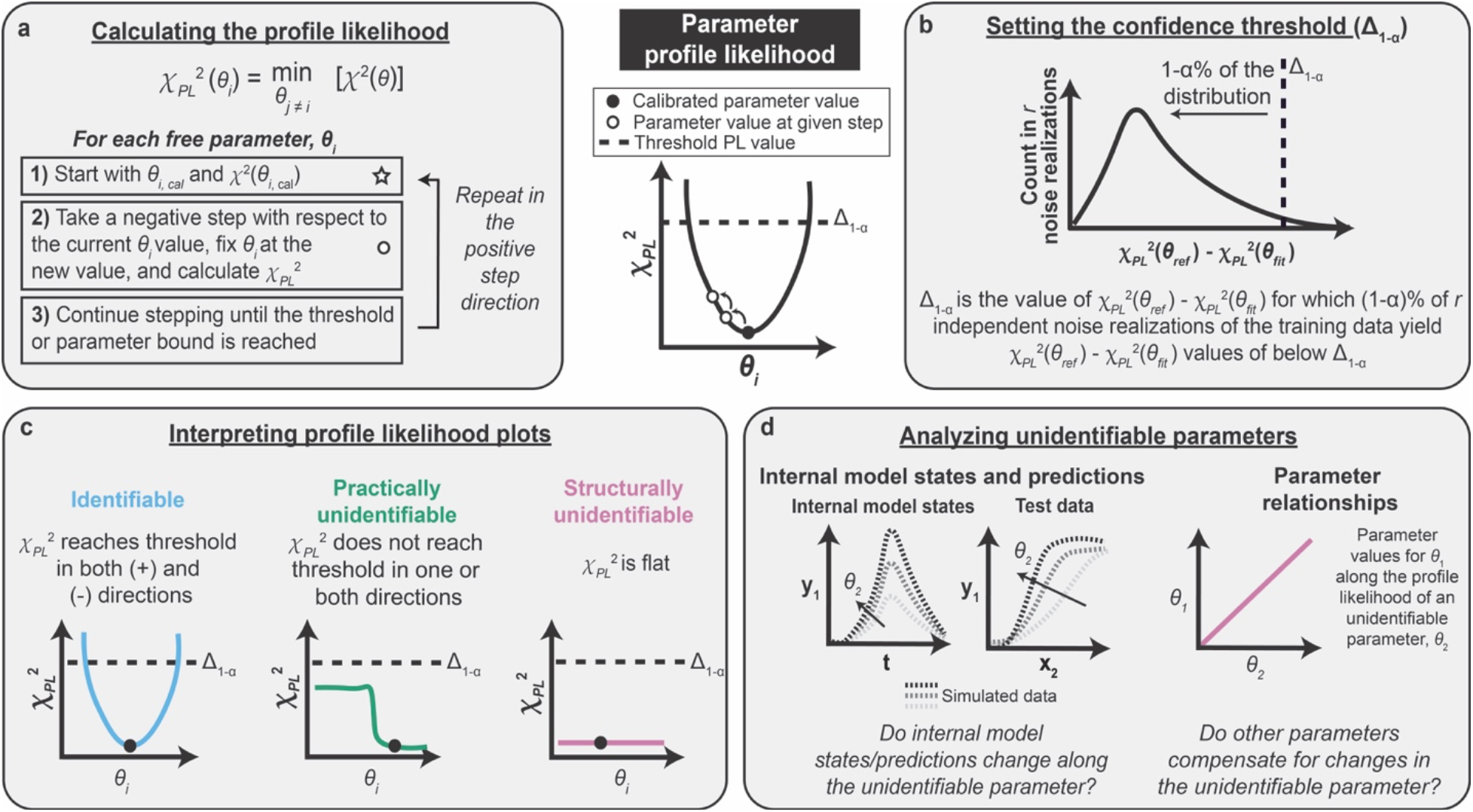
Steps in the parameter profile likelihood approach. **(a)** Calculating the PPL involves iterations of parameter estimation for each free parameter. *θ* is the set of free parameters and *θ_i_* is an individual free parameter. **(b)** The confidence threshold determines the range of parameter values supported by the training data based on the measurement noise. **(c)** The shape of the PPL determines whether a parameter is identifiable, structurally unidentifiable, or practically unidentifiable. **(d)** Unidentifiable parameters can be investigated by analyzing parameter relationships and internal model states and predictions.

- The profile likelihood *χ_PL_*^2^ is determined individually for each parameter by systematically varying the parameter value *θ*_i_, re-optimizing all other parameter values *θ_j≠i_*, and determining the lowest possible value of the cost function χ^*2*^ (**Figure B1a**). The choice of step for each new PPL calculation (each of which is represented by an open circle in the figure) should be chosen such that the step is small when the PPL is steep and large when the PPL is flat. Using an adaptive stepping method facilitates appropriate step sizes and improves computational efficiency (**Supplementary Information**). The PPL approach assumes that the starting point for PPL calculations (the value of the calibrated parameter) resides at a global minimum.
- The confidence threshold Δ_1-*α*_ enables determination of the range of parameter values supported by the experiment by accounting for measurement noise associated with the training data (**Figure B1b**). Parameter sets yielding *χ_PL_^2^* values below Δ_1-*α*_ are considered not significantly different from one another in terms of agreement with the training data. Instead, any improvement upon the PPL below the confidence threshold is due to fitting the noise in the training data and is not indicative of an improvement in the description of the physical system. The choice of threshold is determined by a simulation study in which a number *r* of noise realizations of the training data are generated, representing the variance in training data values that would be expected if the experiment were repeated *r* times (**Supplementary Information**). Each noise realization is defined by adding noise to each datapoint in the original training data set, such that each noise realization is consistent with the original training data but is quantitatively different. For each noise realization, *χ*^2^(*θ_ref_*) is determined by calculating the cost function associated with the noise realization (training data) and the simulated data using the set of reference parameters. *χ*^2^(*θ_fit_*) is determined for each data set by re-optimizing all free parameters with respect to the data set associated with the noise realization. The difference between these values, *χ*^2^(*θ_ref_*) – *χ*^2^ (*θ_fit_*), represents the amount of overfitting for each data set and can be plotted as a histogram for all noise realizations. Δ_1-*α*_ is chosen such that 1-α% of the *χ*^2^(*θ_ref_*) – *χ*^2^(*θ_fit_*) values are below Δ_1-*α*_. In our case study, we use α =0.01. In practical cases in which *θ_ref_* is not known, *θ_ref_* can be approximated as *θ_calibrated_*, as long as the calibrated parameter set yields good agreement with the training data.
- The shape of the profile likelihood determines whether the parameter is structurally unidentifiable, practically unidentifiable, or identifiable (**Figure B1c**). Identifiable parameters do not need refinement, but each unidentifiable parameter should be individually investigated and refined until all parameters are identifiable.
- Unidentifiable parameters can be investigated in two ways (**Figure B1d**). First, the relationships between each unidentifiable parameter and the other parameters can be analyzed. By re-optimizing all other parameter values along the given parameter, the PPL approach enables identification of relationships between parameters. For example, a change in one parameter might be compensated for by a change in another parameter without changing the value of the cost function. The identification of such relationships is an advantage of the PPL approach over other parameter analysis methods, such as a traditional sensitivity analysis where the effect of changing only an individual parameter is evaluated. Second, internal model states and model predictions can be evaluated along each unidentifiable parameter. The impact of each unidentifiable parameter on model predictions and other conclusions can be examined and used to make downstream model reduction and experimental design decisions. For example, if a parameter is unidentifiable and changing the parameter does not change the model predictions, then the parameter can be set to an arbitrary value and the modeler can move on. However, if a parameter is unidentifiable and the model predictions change depending on the value of the parameter, then new training data are required to constrain the unidentifiable parameter. Based on the shape of the PPL (e.g., structurally or practically unidentifiable), directionality of the unidentifiability (e.g., in the positive direction, negative direction, or both), and other factors, there is exists a process that has been described for refining each unidentifiable parameter^*51*^. We note that parameter relationships should be investigated for all unidentifiable parameters to gain information about the sources of unidentifiability before choosing strategies for model reduction or experimental design.

**Figure 7.**
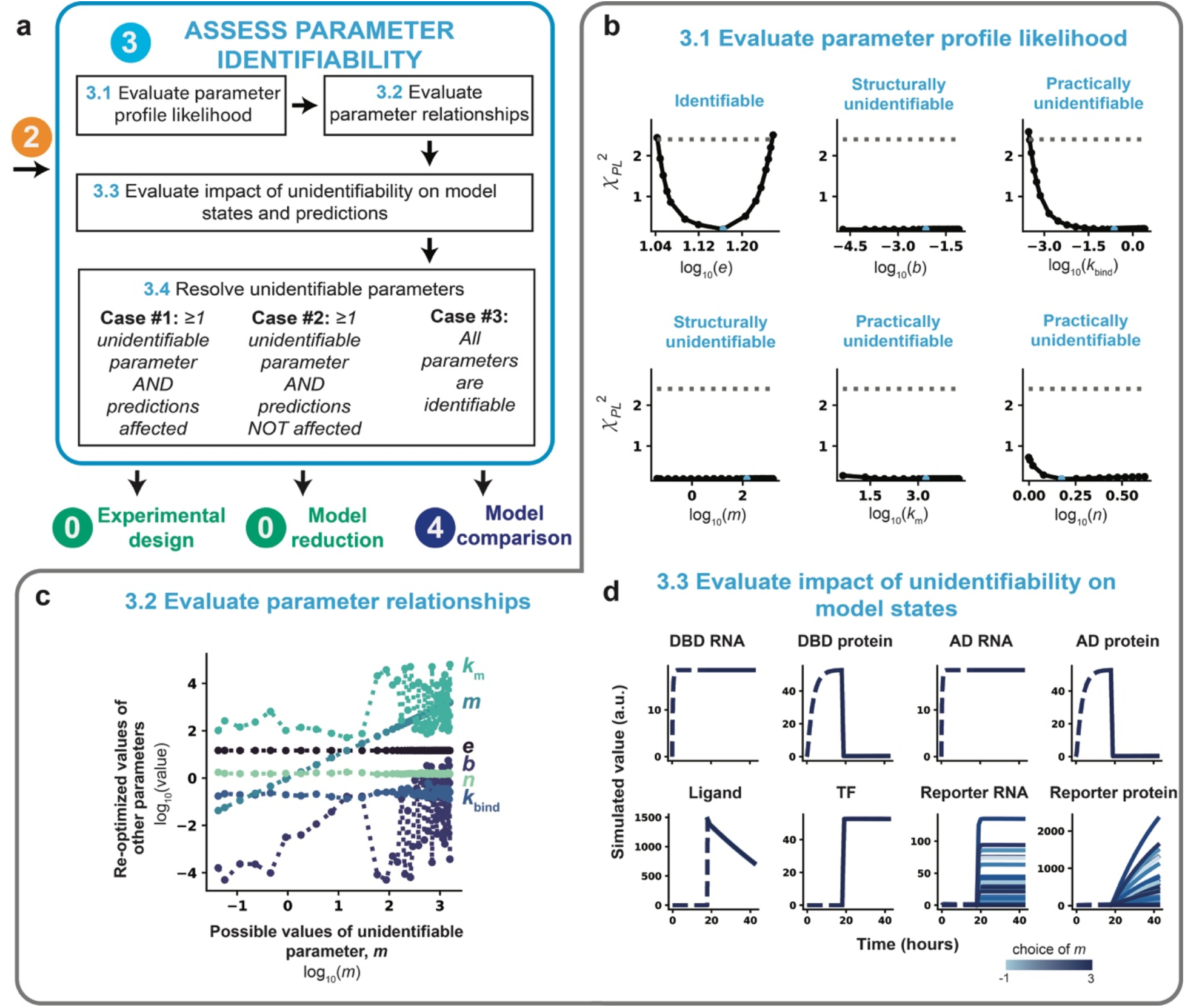
Assess parameter identifiability. **(a)** Module 3 workflow for evaluating and refining parameter identifiability through the profile likelihood approach. Depending on the results of the parameter identifiability analysis, the next step is either experimental design (Module 0), model reduction (Module 0), or model comparison (Module 4). **(b–d)** Module 3 case study for a hypothetical crTF. **(b)** Application of the profile likelihood approach to the model defined in **Figure 2**. The calibrated parameter set from a parameter estimation run with 1000 global search parameter sets, and 100 initial guesses for optimization were used as the starting point (represented in blue). Parameters were allowed to vary across three orders of magnitude in either direction of the reference parameter value, except for *n*, which was allowed to vary across [10^0^, 10^0.6^]. An adaptive step method (**Supplementary Note 1**) was used to determine each step size. The threshold is defined as the 99% confidence interval of the practical *χ^2^_df_* distribution (Δ_1-α_ = 2.4). **(c)** Plots of parameter relationships along the profile likelihood associated with *k*_m_. We consider a range of possible values of the unidentifiable parameter (*m*) and plot these values against recalibrated values of other model parameters (*k*_m_, *e*, *n*, *b*, *k*_bind_). **(d)** Plots of internal model states considering a range of possible values of unidentifiable parameter *m*. Time courses represent the trajectory of each state variable in the model as a function of *m* choice. Each trajectory was generated by holding *m* constant at the given value and reoptimizing all other free parameters (results are from the same simulations used to plot the PPL results in b). Data are shown for these conditions: 50 ng DNA-binding domain plasmid, 50 ng activation domain plasmid, and a saturating ligand dose (100 nM).

In general, structural unidentifiability refers to inherent unidentifiability arising from the model structure itself, while practical unidentifiability captures additional identifiability issues arising from limited or noisy experimental measurements^*51*^. In practice, it is often difficult to distinguish between these two types of unidentifiability because issues with data quality or structural issues (or both) can lead to practical unidentifiability. Flat PPLs are unambiguous indicators of structural unidentifiability, but parameters classified as practically unidentifiable by their PPLs can still be affected by structural issues. For example, if a parameter has a PPL that crosses the threshold in only one direction, the parameter would be classified as practically unidentifiable, but it is possible that the PPL does not cross the threshold in the other direction because of a structural issue, such as a reaction timescale that is much faster than others in the model. Therefore, while the classification of parameters as practically or structurally unidentifiable can be helpful, it is also important for the modeler to determine the cause of each unidentifiability by exploring parameter relationships along each PPL.

As an a posteriori method, the PPL assesses local identifiability of parameters because it infers structural identifiability based on the model fits to the training data, as opposed to a priori methods, which perform the analysis using the equations only and assess global structural unidentifiability^*49*^. Regardless of this limitation, the PPL is an appropriate general method for assessing practical and structural identifiability simultaneously, can be visually interpreted, and provides information that can guide model refinement to improve unidentifiability. Other methods for assessing parameter identifiability are reviewed elsewhere^*37, 51*^. For large models, it may be necessary to use a more computationally efficient algorithm or method to assess parameter unidentifiability^*30, 52, 53*^. Potential failure modes of the PPL approach and suggested remediation strategies are detailed in **Figure S4**.

#### Case study

We executed Module 3 and observed that the base case training data set did not enable identification of most of the free parameters. Using the calibrated parameters chosen in Module 2, we evaluated the PPL individually for each of the six free parameters using a 99% confidence threshold (**Figure 7b**). The threshold was determined to be 2.4 for this model and set of training data based on a simulation study (**Figure S5**, **Supplementary Note 1**). The results demonstrate that only one parameter, *e*, is identifiable given this model structure and training data. Based on the PPL results, all other parameters appear to be either locally practically unidentifiable (*k*_bind_, *n*, *k*_m_) or locally structurally unidentifiable (*b*, *m*). The PPL for *k*_bind_ reaches the threshold in the negative direction, but this parameter is unidentifiable in the positive direction, while the PPLs for the other practically unidentifiable parameters, *n* and *k*_m_, do not reach the threshold in either the negative or positive directions, but are not flat. The PPLs for *b* and *m* are flat, suggesting that these parameters are structurally unidentifiable.

To investigate the relationships underlying each unidentifiable parameter, we first plotted the parameter trajectories along the profile likelihood associated with *m* (**Figure 7c**). The results show either flat (*e*, *n*), nearly flat (*k*_bind_), or bumpy (*b*, *k*_m_) relationships between all parameters. The flat relationships between *m* and the parameters *e* and *n* indicate that these parameters do not compensate for changes in *m* and therefore do not contribute to the unidentifiability of *m*. The nearly flat relationship between *m* and *k*_bind_ indicates that *k*_bind_ does not compensate for changes in *m*; this relationship might be impacted by noise, causing the relationship not to be exactly flat.

The bumpy, seemingly correlated relationships between *m* and the parameters *b* and *k*_m_ suggest that each of these parameters might compensate for one another. In other words, there might be many sets of *b*, *m*, and *k*_m_ values that yield similar fits to the training data, such that it would not be possible, given the model structure and training data, to uniquely identify these parameters. If only two parameters directly compensate for one another, the resulting relationship between the parameters is expected to be smooth, as changes in one parameter are compensated for by another parameter. The complex relationship between *m* and the parameters *b* and *k*_m_ make it difficult to assess exactly how these three parameters depend on one another from a two-dimensional plot. Indeed, when we plotted the values of *m*, *b*, and *k*_m_ along the unidentifiability associated with *m* in three dimensions using the same data generated when evaluating the PPL (**Figure 7a, b**), the surface was smooth. This result indicates that these parameters compensate for one another as *m* is changed and no other parameters are involved in the unidentifiability (**Figure S5b**). We note that parameter relationships in more than two dimensions cannot always be unambiguously recovered using this approach, as only one parameter is fixed at a constant value^*30*^.

Next, to further analyze the consequences of the unidentifiability of *m*, we plotted the internal model states along the profile likelihood associated with *m* (the set of fixed *m* values and other re-optimized parameter values considered equally plausible based upon the analysis in Figure 7b) at saturating ligand dose (**Figure 7d**). All states upstream of promoter activation are indistinguishable along the profile likelihood (model states shown in blue for a scale of fixed *m* values). The dynamic trajectories for both the reporter RNA and reporter protein states along the profile likelihood of *m* are similar in shape but differ in scale. Therefore, the unidentifiability of *m* does affect the dynamic trajectories of the internal model states. We note that the profile likelihood of *m* may also be related to the data-driven normalization scheme used in which both the training data set and each simulated data set are both divided by their respective maximum values before comparison and no other readouts are measured in units that are comparable to the reporter protein. Although not investigated here, we note that the normalization strategy should also be explored and investigated in a context-specific way when refining parameter identifiability (**Supplementary Table 1**). Normalization can be thought of as a feature of the model structure; if parameters are not identifiable, one should consider changing the normalization based on intuition and evaluating that change via the PPL.

### Refinement of parameter identifiability using experimental design and model reduction

#### Motivation

If the parameters are not identifiable given the training data, then the model is considered overly complex, and there should exist a simpler model that explains the data equally well. Alternatively, additional training data could be collected to further constrain parameter estimates. The choice of model reduction or additional data collection depends on whether simplifying the model reduces the explanatory or predictive power of the model. Specifically, if an unidentifiable parameter does not affect predictions, then the model can be reduced to remove the parameter, but if an unidentifiable parameter does affect predictions, then additional training data should be incorporated to constrain the parameter and therefore the predictions^*31*^. Multiple rounds of refinement may be necessary to arrive at a fully identifiable model.

#### General method

After each parameter is classified based on the PPL, one can use model reduction and experimental design to refine the model and improve identifiability (**Figure 7a Module 3.4**). The type of unidentifiability and the shape of the PPL are used to determine a promising model refinement strategy. We consider three cases outlined in **Figure 7a** to illustrate this approach. In Case 1, if at least one parameter is unidentifiable and model predictions are impacted, additional training data are proposed and evaluated via proposition, collection, and incorporation of these data in Module 0. This step can include collecting experimental data with reduced noise, with additional independent variables or observables, or with different sampling procedures. In Case 2, if at least one parameter is unidentifiable, but model predictions are not impacted, additional training data are not necessary, and the model is instead reduced by returning to Module 0 and proposing a reduced model (i.e., a revised set of ODEs that describe an appropriately simplified mechanism). If the parameter is structurally unidentifiable, the parameter can be fixed to a constant value^*34*^. If the parameter is practically unidentifiable, a model reduction strategy based on the shape of the PPL (e.g., whether there is a bound in the positive direction, negative direction, or both) can be pursued^*34*^. In cases for which the modeler cannot use intuition to choose a model reduction strategy, semi-automated model reduction strategies can identify parameter combinations and suggest how to remove unnecessary model terms and variables by taking asymptotic limits^*54*^. In Case 3, all parameters are identifiable, and the modeler proceeds to Module 4 in which competing candidate models are compared to one another. Through systematic iteration between models and experiments, the PPL enables the modeler to assess whether the final model is appropriately complex given the available data.

#### Case study

We hypothesized that incorporating additional information about how the plasmid doses of the DNA-binding domain and activation domain components impact reporter expression would resolve the unidentifiability of *b*, *m*, *k*_bind_, *n*, and *k*_m_. To test this hypothesis, we generated additional training data (using our reference model, for the purposes of this tutorial) varying the dose of plasmid encoding DNA-binding domain in combination with two doses of plasmid encoding activation domain (20 ng, 10 ng) and the saturating ligand dose of 100 nM (**Figure 8a**). The additional training data were added to the original ligand dose response data set (Model A) to define Model B. The dependence of reporter expression on DNA-binding domain plasmid dose is non-monotonic, which is physically plausible as excess DNA-binding domain (at high DNA-binding domain plasmid doses) acts as an inhibitor of promoter activation. Before evaluating the PPL, the parameter estimation method chosen in Module 1 was evaluated under the new conditions (including the additional training data), hyperparameters were tuned, and a new calibrated parameter set with an R^2^ of 0.999 was identified (**Figure S6**). The new calibrated parameter set was used to initialize simulations for the PPL (**Figure 8b**).

**Figure 8.**
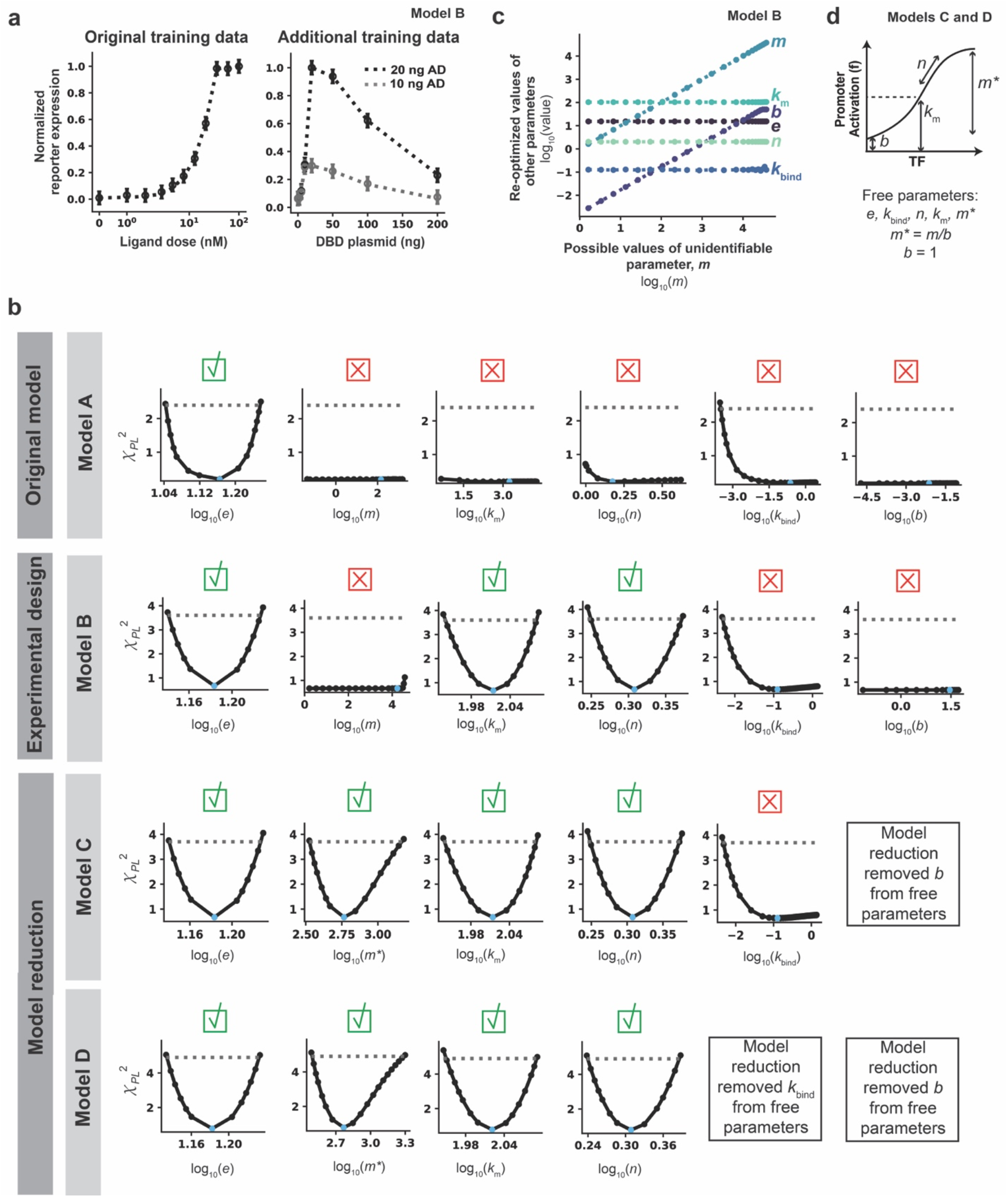
Refinement of parameter identifiability using experimental design and model reduction. **(a)** Model B training data. Additional training data for a DNA-binding domain plasmid dose response at two plasmid doses of activation domain and a saturating ligand dose were generated using the reference parameter set. Noise was added as was done for the ligand dose response. Model B has the same model structure as Model A, but Model B incorporates this additional training data set and therefore has different calibrated parameters. **(b)** PPL results for Models A, B, C, and D. Results from Model A (Figure 7b) are shown again to facilitate comparison between other PPLs for other models. Each column is a different parameter, and each row is a different model. All parameters were allowed to vary across three orders of magnitude in either direction of the reference parameter value, except for *n*, which was allowed to vary across [10^0^, 10^0.6^] for all PPL simulations in this figure. Calibrated parameter values for Models B, C, and D are in **Supplementary Table 2**. The threshold is defined as the 99% confidence interval of the *χ_df_^2^* distribution (Model B: Δ_1-*α*_ = 3.6, Model C: Δ_1-*α*_ = 3.7, Model D: Δ_1-*α*_ = 4.9). A green check mark means the parameter is identifiable and a red X means the parameter is unidentifiable. **(c)** Parameter relationships along the profile likelihood associated with *m*. *b* and *m* compensate for one another along the profile likelihood. **(d)** Model reduction scheme for Model C. Instead of fitting both *m* and *b*, the ratio between the two parameters was fit and *b* was fixed to an arbitrary value of 1.

Incorporating the additional training data enabled the identification of two previously unidentifiable parameters (*k*m and *n*), and the remaining parameters appear either locally structurally unidentifiable (*b*) or locally practically unidentifiable (*m* and *k*_bind_). While the PPL for *m* with Model A indicated that *m* was structurally unidentifiable, the PPL for *m* with Model B indicated that the parameter is practically unidentifiable, with an increase in *χ_PL_^2^* at high values of *m;* these high values of *m* were not explored in the PPL for Model A because the calibrated value of *m* was much lower in Model A, and therefore the maximum number of PPL steps was reached before such a high fixed *m* value was reached. To investigate this inconsistency, we plotted parameter relationships along the profile likelihood associated with *m* (**Figure 8c**). For most values of *m* considered, the value of *b* depended on the value of *m*, indicating that the values of these parameters can compensate for one another without changing the simulated training data. *k*_bind_ is still practically unidentifiable, but this parameter is not involved in the structural unidentifiability of *b* and *m*, as the relationship between *m* and *k*_bind_ is nearly flat. Now that only two parameters are involved in the unidentifiability, a two-dimensional visualization shows the relationship between *b* and *m* in a clear way that was not possible in the parameter relationship plots for Model A. At very high values of *m* (above 10^4^), *b* remains constant at its maximum bound and can no longer increase to compensate for increases in *m*. This result shows the cause of the unidentifiability and resulting dependencies of *m* and *b*.

These results illustrate a limitation in that PPL evaluates local identifiability rather than global identifiability, leading to inconsistent PPL classifications across models. The results support the need to consider both parts of the identifiability analysis (PPL plots and parameter relationships) to investigate unidentifiable parameters and make model refinement decisions. For our case study, we used this investigation to justify the inconsistent PPL classifications. If inconsistencies cannot be justified by the suggested investigations, such as for larger, complex models in which many parameter values compensate for one another, the modeler may need to incorporate a global, a priori method for parameter identifiability analysis^*37, 51*^. Overall, Model B is an improvement over Model A in terms of the number of identifiable parameters, but it did not meet the requirement for all parameters to be identifiable. This case study illustrates the common experience that more than one round of refinement may be necessary to arrive at a final model.

To resolve the unidentifiability related to *b* and *m*, we reduce the model by estimating the relative magnitude of *b* and *m* rather than each value separately (**Figure 8d**). The value of *b* was arbitrarily set to 1, and we introduce and fit a new parameter, *m*\*, that describes the fold induction of promoter activation (maximum value at a saturating amount of crTF divided by background value with no crTF). This substitution reduces the total number of free parameters from six in Models A and B to five in Model C. Before evaluating the PPL, the parameter estimation method was evaluated under the new conditions (new model structure), hyperparameters were tuned, and a new calibrated parameter set with an overall R^2^ of 0.999 was identified (**Figure S7**). The new calibrated parameter set was used to initialize simulations for the parameter profile likelihood (**Figure 8c**). The model reduction scheme in Model C obviates the need to constrain *b* (which is now a fixed parameter) and enables identification of *m*\*. Now the only remaining unidentifiable parameter is *k*_bind_, which is practically unidentifiable.

To resolve the unidentifiability of *k*_bind_, we defined Model D and incorporated a simple model reduction in which *k*_bind_ was arbitrarily set to 1. This choice, similar to the others in this section, is one of many appropriate strategies for model reduction or experimental design that could be pursued. For example, we could propose additional training data with which to constrain *k*_bind_ rather than setting it to an arbitrary value; this would be the preferable option if there were model predictions affected by the unidentifiability of *k*_bind_. The PPL results, in which *k*_bind_ has a bound in the negative direction but not in the positive direction, indicate that *k*_bind_ can take any value as long as it is sufficiently large, representing fast binding between crTF components. As long as the rate constant for binding is large enough, the components will bind very quickly, and no increase in this rate will affect the model agreement with the training data. Although *k*_bind_ is classified as practically unidentifiable by the PPL, the results indicate that the cause of this unidentifiability is structural and can be attributed to a separation of timescales between the crTF component binding reaction and other reactions in the model. Therefore, in this case study, fixing *k*_bind_ to an arbitrarily large value is appropriate. We note that the utility of a model reduction strategy can be assessed by comparison of information criteria that take into account both the fit to training data and the number of free parameters^*26*^. Given a set of candidate models, comparison of information criteria can determine which model most likely matches the data; this approach is addressed in Module 4.

Before evaluating the PPL for Model D, the parameter estimation method was evaluated under the new conditions (new set of free parameters), hyperparameters were tuned, and a new calibrated parameter set with an overall R^2^ = 0.993 was identified (**Figure S8**). The new calibrated parameter set was then used to initialize simulations for the parameter profile likelihood (**Figure 8c**). The PPL indicated that all free parameters were identifiable in Model D, and therefore no further rounds of model reduction or experimental design were necessary.

### Module 4: Compare candidate models

#### Motivation

Different models may yield similar agreement with experimental data. The final challenge is to compare these models and choose the best candidate.

#### General method

Models can be compared based upon the information criterion (**Figure 9a Module 4.2**) and based upon model agreement with training and test data (**Figure 9a, Module 4.1, 5.3**).

**Figure 9.**
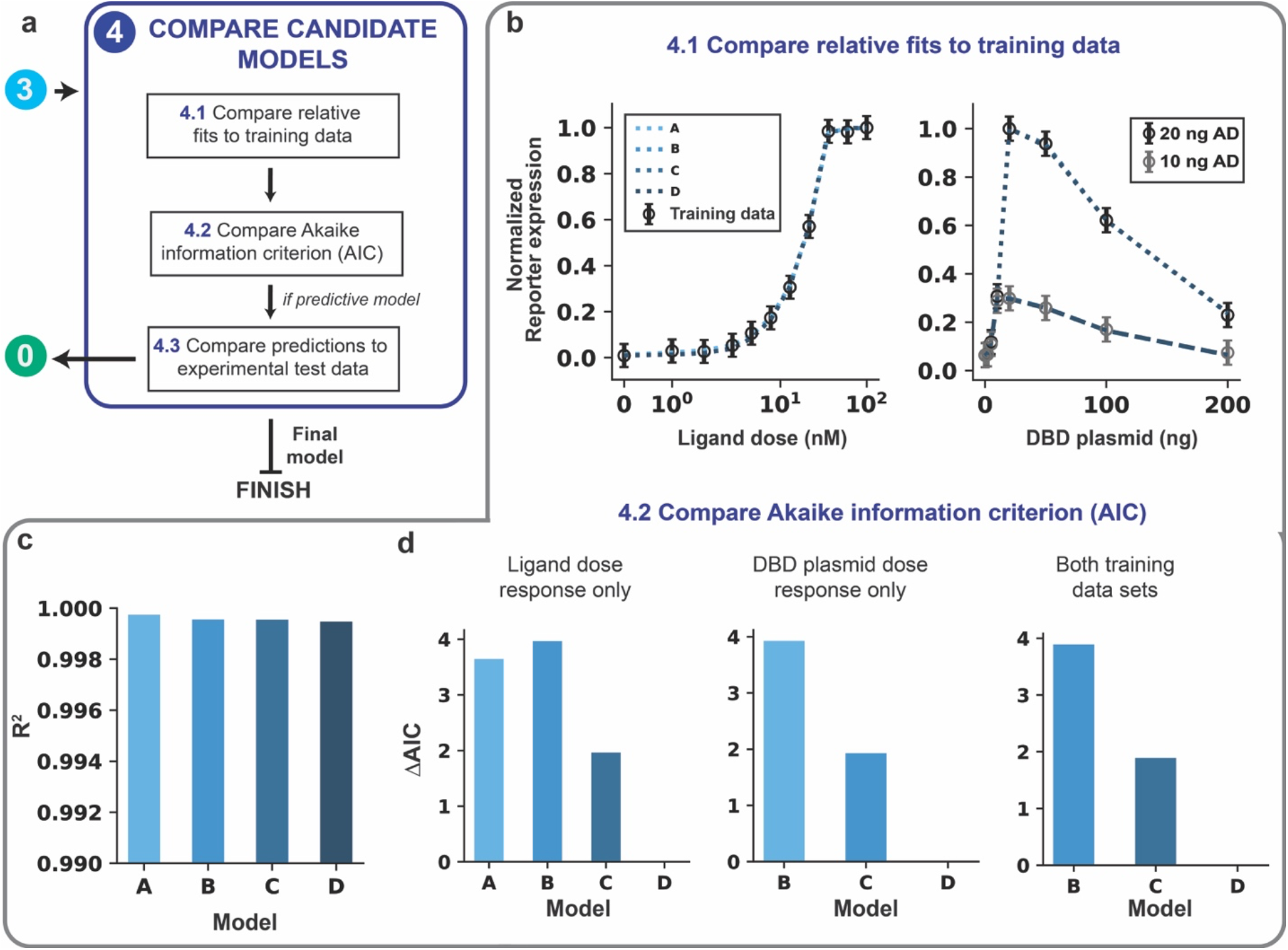
Compare candidate models. **(a)** Module 4 workflow for model comparison. If competing models remain after parameter identifiability analysis, then the models are compared on the fit to training data using both the AIC and comparison of predictions (if the model aims to be predictive). **(b–d)** Module 4 case study for a hypothetical crTF. **(b)** Comparison of the relative fits to training data for Models A, B, C, and D (simulations overlap). **(c)** Comparison of quantitative fit to training data based on R^2^. A high R^2^ (max = 1) is ideal. **(d)** Comparison of AIC differences for using AIC calculated with ligand dose response data only (left), DBD plasmid dose response data only (middle), and both data sets (right).

The Akaike information criterion (AIC)^*26, 27, 55*^ is a metric that is used to quantitatively compare models **(Equation 12)**. The AIC is defined as:

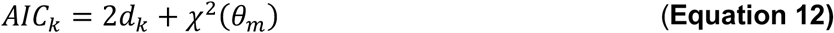

where *d_k_* is the number of free parameters in model *k* with calibrated parameter set *θ_m_*. The AIC ranks models accounting for both the model complexity (first term) and the fit to training data (second term). Lower AIC values indicate greater parsimony. The intent of including a term to penalize complexity is to prioritize models that are generalizable over those that have unnecessary mechanisms or unidentifiable free parameters. Given a set of nested candidate models (with one being a special case of another, as is the case here), the best model chosen based on the information criterion should by definition be the same as the model that is chosen based on the identifiability analysis. However, in cases where candidate models are not nested and multiple models are fully identifiable, the AIC is a useful metric for making comparisons. The AIC must be calculated for each model using the same data. Therefore, if two models were trained with different data sets, the error in fit must be recalculated using a consistent data set to determine the AIC.

To compare relative AIC values between competing models, the AIC difference (Δ*AIC_k_*) for model *k* is calculated as:

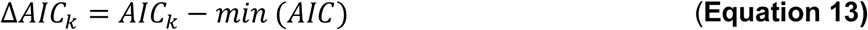

Where (*AIC*) is the minimum AIC value among the candidate models^*26*^. A small AIC difference for a model *k* (Δ*AIC_k_* ≤ 2) indicates substantial support for the model, meaning that both model *k* and the model with the minimum AIC are supported by the information criterion analysis. Moderate differences (2 ≤ Δ*AlC_k_* ≤ 10) indicate less support, and large differences (Δ*AlC_k_>* 10) indicate no support^*56*^. We note that these guidelines do not always hold, especially when experimental observations are not independent.

If upon further experimental investigation (i.e., to generate empirical test data), none of the competing models provide adequate predictions, then the modeler can return to Module 0 and either propose new mechanistic assumptions or collect new training data. If the model predictions cannot be experimentally recapitulated, then it might be necessary to incorporate the original test data into the training data set, generate new candidate models, and test them against new experimental test data.

#### Case study

We use the previously generated Models A, B, C, and D to demonstrate the model comparison process and select the best candidate. The AIC values are compared separately for the two datasets (ligand dose response and DNA-binding domain plasmid dose response) and for the combination of datasets (**Figure 9d**). For the ligand dose response only, the model with the lowest AIC is Model D, while Model C is supported by the information criterion analysis (Δ*AIC~2*), and Models A and B are less supported (**Figure 9d**, left). For the DNA-binding domain plasmid dose response only, the model with the lowest AIC is again Model D, while Model C is again supported by the information criterion analysis, and B is less supported (**Figure 9d**, middle). For both training data sets, only Models B, C, and D (which were each trained with both data sets) can be compared. Similar to the previous results, Model D has the lowest AIC, Model C is again supported by the information criterion analysis (*ΔAIC~2*), and Model B is less supported (**Figure 9d**, right). The results from this module are consistent with the results of the parameter identifiability analysis (Module 3), which indicate that Model D is the best model. Formally, the information criterion analysis does not justify rejection of Model C, but we can justify the choice of Model D over Model C because Model D is the only one with no unidentifiable parameters. Model D is an example of a fully identifiable final model that is the ideal output of GAMES.

### Module failure modes and suggested remediation

Each module has a unique set of failure modes that can be addressed by altering specific workflow choices (**Supplementary Table 3**, **Figure S4**). The case study in this tutorial represents an ideal version of the workflow without any module failures. We note that the GAMES workflow refers to the overall, conceptual workflow used to step through the model development process, not the specific methods used in this example, which may need to be adjusted for different modeling problems. In practice, alterations such as those described in **Supplementary Table 3 and Figure S4** may be necessary to arrive at a model that satisfies a given modeling objective and that has identifiable parameters.

### Iteration between computational tasks and experiments

The model development process is iterative, and different modules can be combined to accomplish separate, complementary goals (**Figure 10**). *Parameter estimation* yields a parameter set or sets that confer the best agreement between the training data (normally, experimental data) and simulations for a given model structure. *Model refinement* includes an option for iteration between modeling and experiments if a good fit is not obtained using the initial model. *Experimental design* employs parameter identifiability analysis to propose and validate the use of additional training data to constrain parameters. *Model reduction* employs parameter identifiability analysis to drive the selection and validation of appropriate strategies to simplify the model. *Model selection* employs the first four modules to produce a set of competing models and then compares the resulting best-case versions. Finally, *model validation* compares simulated test data (predictions) with experimental test data and can involve model refinement to improve this agreement as needed.

**Figure 10.**
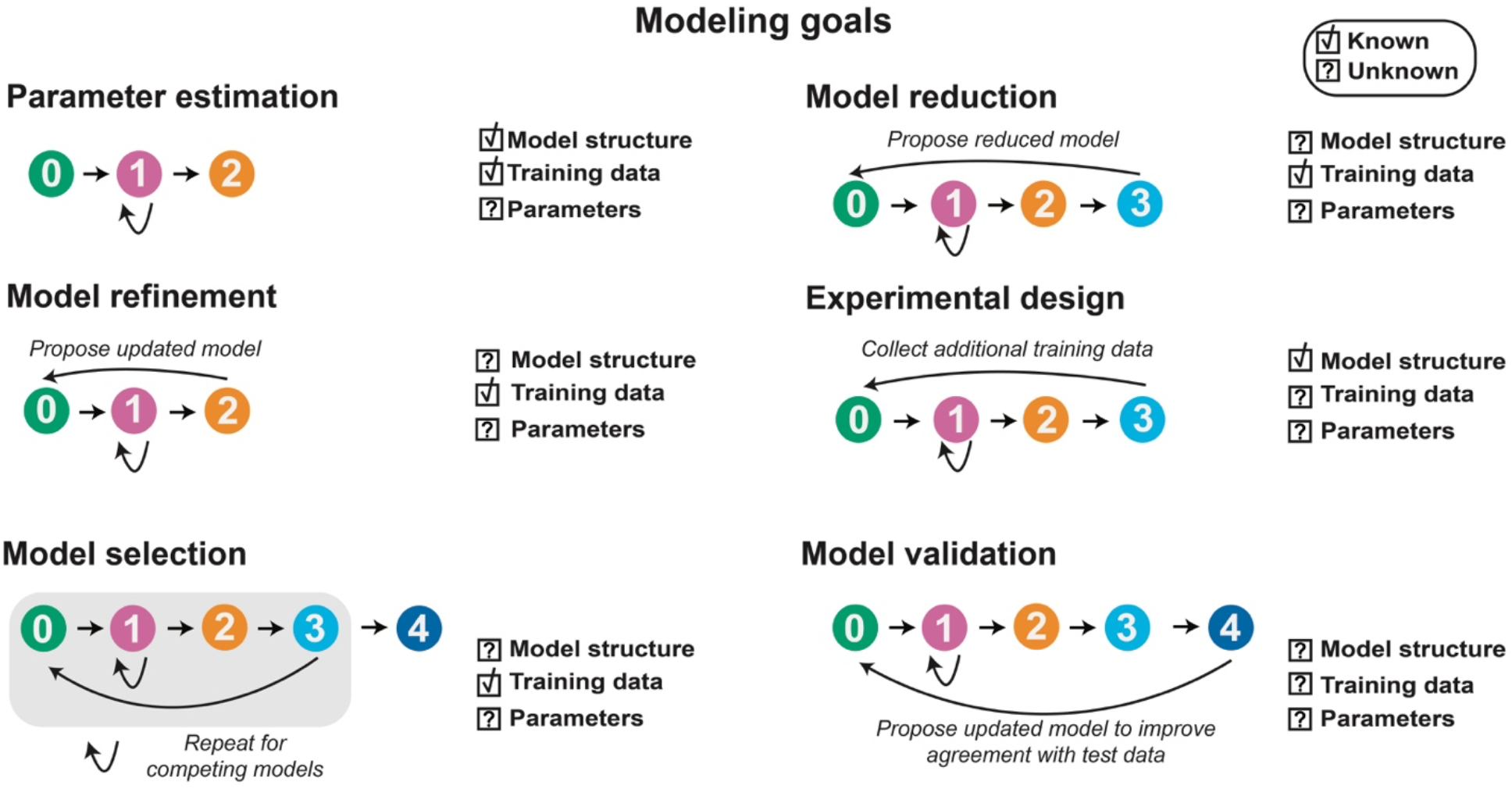
The modeling workflow is iterative and can be deconstructed to accomplish different modeling goals. Model development is deconstructed into different goals. Boxes with check marks indicate that a feature is assumed to be known, and boxes with question marks indicate that a feature will be identified or determined as part of meeting a specified goal. Arrows indicate cycles that are conducted until a condition is met.

## Summary and discussion

GAMES is generalizable and can be adapted for different goals, independent of the specific biological and biochemical parts under investigation. By considering each of the modules described here, researchers can increase the power and utility of their modeling efforts by using the model development process to gain understanding about a system of interest, rather than focusing only on generating a single calibrated model, for which parameter uncertainty and predictive utility may not have been systematically evaluated. The workflow in this tutorial introduces and uses aspects of model development that should improve the rigor and reproducibility of modeling in synthetic biology, which represents an open opportunity and need^*16, 17*^. We also anticipate that this tutorial could be used as an educational tool.

To support access for researchers, educators, and students, the GAMES code (**Methods**) was written in the freely available programming language Python. The code can be assessed on GitHub, enabling reproducibility and reusability. We anticipate that GAMES could be adapted to previously developed software packages such as Data2Dynamics^*57*^, PottersWheel^*58*^, COPASI^*59*^ (a Complex Pathway Simulator), and BMSS^*60, 61*^ (an automated Biomodel Selection System) to accomplish specific computational tasks, and that this adaptation could improve computational efficiency compared to the code in this tutorial. Our code is intended primarily to provide an example of how to implement GAMES. The utility of starting from scratch without relying on software packages is a greater flexibility for customization. However, there is a tradeoff between customizability and the time and effort to be considered by the modeler, as building a code base from scratch can be time-consuming and prone to error.

GAMES provides a systematic model development process, avoiding brute force consideration of all possible options, such as model structures, parameter estimation methods, or sets of training data, which may be infeasible in many cases. For example, if a modeler does not know which parameter estimation method is well-suited to the problem of interest, one could test many different methods, without careful consideration of the theory behind each method, until a method that passes the PEM evaluation criterion is identified. However, this approach is not encouraged. Instead, we recommend that the modeler use this workflow to test hypotheses as to whether or not a given method is suitable for a modeling objective. The modeler should formulate a hypothesis as to why a given method is appropriate for an objective and then use the workflow to test that hypothesis, for example, “We hypothesize that the cost function has multiple minima and therefore a multi-start Levenberg Marquardt algorithm with the following specified hyperparameters would be appropriate for balancing global parameter exploration and speed for finding local minima.” GAMES enables the modeler to evaluate whether this hypothesis holds and either tune hyperparameters or try a different algorithm if it does not. Similarly, this principle can be applied to model formulation. One could imagine comparing a large, somewhat randomly determined set of models without considering the biological relevance of each one. However, a preferred path is for the modeler to formulate a hypothesis as to how the system works, such as “Even though true trimolecular interactions are rare in nature, we hypothesize that the interaction of these three components can be suitably described by a single, irreversible, trimolecular reaction,” and then rigorously test that hypothesis using GAMES. This approach is different than considering all possible combinations of reactions that could potentially occur, which has additional training data requirements and has become possible only recently by using specialized sparse-optimization methods^*62–64*^.

The field of synthetic biology is well-positioned to utilize ODE models due to the wide range of potential experiments that can be designed to interrogate or perturb synthetic systems. There are often many tunable handles, including experimental conditions (e.g., amounts of each component in a genetic program), design choices (e.g., substitution of parts with the same functionality but different quantitative performance), and topologies (different interactions between components). Performing comprehensive experiments to investigate the effects of each tunable handle, individually and in combination with each other handle, is infeasible. For this reason, ODE models have proven invaluable by decreasing the number of experiments needed to better understand or predict the response of a genetic program to these perturbations^*5, 7*^. GAMES enables these investigations by providing an accessible, systematic, and reproducible method to ensure that a set of experimental observations and model structure are together appropriate to generate useful predictions.

We anticipate that the core GAMES framework may be improved and extended in future iterations. Opportunities for improvement include developing user-friendly software based on the example code and incorporating more method options for specific types of problems, such as parameter estimation methods designed to handle noisy data^*23*^ or to improve PEM efficiency when parameter unidentifiability leads to nonconvergence^*45*^. Opportunities for extension include: (1) utilizing the method in this tutorial to build, analyze, and employ other synthetic biology systems; (2) improving predictive design; and (3) integrating automated computational design methods and human decisionmaking (e.g., using GUIs or automation of processes such as model selection^*62–64*^ or in silico experimental design). We hope that continually improving the accessibility and rigor of computational modeling will facilitate the ongoing evolution of synthetic biology as a technical discipline.

## Methods

### Approximation of dynamics for case study

Custom Python scripts (Python 3.7.3) and Python’s odeint solver were used to run simulations. Equations were solved using two sequential simulations analogous to the experimental procedure associated with transient transfection, a common DNA delivery method used to prototype genetic programs^*5, 6*^. The first simulation modeled the system trajectory from the time of transfection (when plasmids for the DNA-binding domain, activation domain, and reporter are delivered to cells) until the time of ligand treatment (18 h). The final time point concentration of each state from the first simulation was then used to initialize a second simulation that modeled the system trajectory from the time of ligand treatment to the time of fluorescence quantification by flow cytometry (24 h). Transient transfection delivers varied plasmid amounts to cells in a cell population, and although this population heterogeneity can be described with a statistical model^*6*^, for simplicity, consideration of population heterogeneity was omitted in this tutorial and the simulations.

### Normalization strategy

Data-driven normalization was used to make comparisons between training data (representative of the process one usually employs for experimental data) and simulations. For each experiment, each data point was divided by the maximum value in the experiment. The same normalization strategy was applied to each simulated data set such that each data point was divided by the maximum value in the simulated data set. In this manner, independent experiments can be qualitatively but not quantitatively compared. In the case study, we assume that the ligand dose response and the DNA-binding domain plasmid dose response were completed in separate experiments and therefore each data set is independently normalized. The effect of normalization procedures on model development and parameter estimation has been examined in another study^*24*^.

### Error metrics

The standard error of the mean (SEM) encompasses biological and technical variation. The standard deviation for each simulated datapoint was set to a constant, arbitrary value of 0.05 arbitrary units (∪). The SEM is calculated based on the standard deviation of measurements, which here are in triplicate (*n_replicates_* = 3):

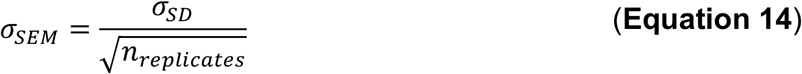

### Parameter estimation method

A multi-start local optimization algorithm was used to estimate parameters (**Figure S3**). The algorithm was implemented using custom Python scripts along with Python packages SALib^*65*^ for global search and LMFit^*66*^ for optimization.

### Parameter profile likelihood

All confidence intervals in this study are *α* = 0.01. Custom Python scripts were used to calculate the PPL based on the algorithm described previously^*30, 37*^. Further explanation is in **Supplementary Note 1**.

### Parallelization of computational tasks

Simulations were parallelized across eight independent cores (chosen based on the number of cores available in the hardware used to run the simulations) to improve computational efficiency. Custom Python scripts were used to implement parallelization. An analysis of the effect of parallelization on computational time has been conducted in another study^*67*^.

## Code availability

Code is provided at https://github.com/leonardlab/GAMES under an open-source license. Simulation outputs are provided as **Supplementary Data 1**.

## Author Contributions

Conceptualization: K.E.D., J.N.L.; Methodology: K.E.D., J.J.M; Writing – Original draft: K.E.D. Writing – Editing and final draft: K.E.D., J.J.M., N.M., N.B., J.N.L.; Visualization: K.E.D.; Software: K.E.D.; Supervision: N.B., J.N.L.; Project Administration: N.B., J.N.L.; Funding Acquisition: K.E.D., N.B., J.N.L.

## Acknowledgements

We thank Austin Chen, Kathleen Dreyer, Jithin George, Sasha Shirman, Alexis Prybutok, Jessica Yu, and members of the Leonard Lab and Bagheri Lab for helpful discussions. This work was supported in part by the National Science Foundation Graduate Research Fellowship Program (DGE-1842165: K.E.D.) and the National Institute of Biomedical Imaging and Bioengineering of the NIH (1R01EB026510: J.N.L.).

