## Supplementary Information for "GAMES: A dynamic model development workflow for rigorous characterization of synthetic genetic systems"

**This file includes:**

Abbreviations

Glossary of terms and definitions

Supplementary Note 1: Practical considerations for parameter identifiability analysis

Supplementary Note 2: Description of functions included in source code and suggestions for adapting the code to new models

Supplementary Note 3: Description of simulation outputs and source data

Figures S1–S8

Supplementary Tables 1–3

References cited in Supplementary Material

### Abbreviations

AD: activation domain
AIC: Akaike information criterion
crTF: chemically responsive transcription factor
DBD: DNA-binding domain
GAMES: Generation and Analysis of Models for Exploring Synthetic systems
L-M: Levenberg-Marquardt
PIA: parameter identifiability analysis
PEM: parameter estimation method
PPL: parameter profile likelihood
TF: transcription factor

### Glossary of terms and definitions

***Calibrated parameter set:*** The parameter set yielding the lowest value for a cost function after parameter estimation; the highest-performing parameter set

***Cost function****:* A metric used to quantitatively compare simulation values and experimental training data during parameter estimation

***Consistency check****:* A step in the parameter estimation process that aims to ensure that the parameter estimation method is properly implemented and that the parameter estimation method, covariates, and hyperparameters are appropriate for the given problem

***Covariate:*** A choice that is not of primary interest but can impact the interpretation of results, e.g., ODE solver, parameter estimation method

***Explanation****:* A modeling objective in which the goal is to help a user describe and understand a set of experimental observations

***Free parameters:*** Parameters in the model that are directly fit using training data

***Modeling scope****:* A modeling decision distinguishing which parts of the system will be described, and which will be omitted

***Hyperparameters:*** Variables that impact the parameter estimation method but are not involved in the model itself; hyperparameters are not kinetic parameters

***Identifiable:*** A property of a parameter that means that the parameter can be uniquely estimated within a finite confidence interval

***Parameter estimation method (PEM):*** A specific algorithm to accomplish parameter estimation; there are many parameter estimation methods that can be applied in different scenarios

***Parameter estimation problem:*** A specific model structure, set of training data, and cost function for which parameter estimation is executed

***PEM evaluation data:*** Simulated data sets generated to evaluate the PEM and tune hyperparameters in Module 1 only; these data sets are generated with the same model structure as defined in Module 0 with sets of arbitrarily chosen PEM evaluation parameters

***PEM evaluation parameters:*** Parameters chosen for PEM evaluation in Module 1 only; these parameters are chosen to yield PEM evaluation datasets that are qualitatively similar to the training data, when possible

***Prediction**: A modeling objective in which the goal is to simulate the response of a system to a previously untested experimental condition or design choice

***Test data:*** A set of observations that a model is intended to predict; test data are not used to estimate parameters

***Training data:*** A set of observations directly used to estimate parameters

***Reference model:*** The model structure generated for the purpose of the case study

***Reference parameters:*** The parameters arbitrarily chosen to define the reference model for the purpose of the case study only

### Supplementary Note 1: Practical considerations for parameter identifiability analysis

#### Determination of the confidence threshold

The confidence threshold $\Delta_{1-\alpha}$ can be determined via likelihood-based confidence intervals as described previously[1, 2]. The confidence threshold is defined as the 1–$\alpha$ quantile of the $\chi^2$ distribution such that:

$$\Delta_{\boldsymbol{\alpha}} = Q({\chi^2}_{df}, 1-\alpha) \qquad \textbf{(Equation S1)}$$

Where ${\chi^2}_{df}$ is the $\chi^2$ distribution corresponding to the number of free parameters as defined by df, df is the number of free parameters in the system, representing a simultaneous confidence interval (instead of a pointwise confidence interval, in which df = 1 and changes to only a single parameter are considered), and $1-\alpha$ is the confidence level (for a 99% confidence level, $\alpha = 0.01$). The $\chi^2$ distribution represents the amount of overfitting by quantifying the expected distribution of the difference $\chi^2(\theta_{ref}) - \chi^2(\theta_{fit})$ across many different training datasets that are statistically consistent with the original training dataset, but differ in exact magnitude due to technical error. For each noise realization, $\chi^2(\theta_{ref})$ is determined by calculating the cost function $\chi^2$ associated with the noise realization (training data) and the simulated data using the set of true parameters $\theta_{ref}$. $\chi^2(\theta_{fit})$ is determined for each noise realization by re-optimizing all free parameters with respect to the dataset associated with the noise realization. $\theta_{fit}$ is defined as the resulting set of calibrated parameters. Each training dataset generated in this way can be considered a unique noise realization.

Although $\Delta_{1-\alpha}$ theoretically can be determined using only the confidence level and the number of free parameters, the actual $\chi^2$ distribution associated with a given model structure, training dataset, and measurement error can differ from the theoretical distribution, especially for nonlinear models and small training datasets**. The actual $\chi^2$ distribution can and should be validated through simulation studies for each new model structure or training dataset.** In case studies such as this one, validation of the $\chi^2$ distribution requires knowledge of the reference parameters underlying the system. In practical cases in which the reference parameters are unknown and estimation of these parameters is part of the modeling objective, the calibrated parameters can be used to define a set of reference training data, which can then be used in a simulation study to validate the $\chi^2$ distribution.

We evaluated the $\chi^2$ distribution and determined the confidence threshold for each model (Model A - Figure S4, Model B - Figure S5c, Model C - Figure S6c, Model D - Figure S7c). To evaluate the distribution, the following steps were taken:

1) Using the standard error of each data point, 1000 unique noise realizations of the training data were randomly generated. Each of these new datasets is statistically

in agreement with the original experimental dataset, but none are identical due to unique noise implementations.

2) For each of these 1000 datasets (noise realizations), we calculated $\chi^2(\theta_{true})$ using the true parameters and $\chi^2(\theta_{fit})$ by re-optimizing all free parameters with respect to the given noise realization.

3) The difference $\chi^2(\theta_{ref}) - \chi^2(\theta_{fit})$ was calculated for each of the noise realizations, and the results were plotted as a histogram (**Figure S7**). The x-axis depicts the amount of overfitting. The threshold is determined by calculating the x-axis value $\chi^2(\theta_{ref}) - \chi^2(\theta_{fit})$ such that $(1-\alpha)$% of the the histogram is below the threshold value. In this case study, the threshold was calculated such that 99% of the $\chi^2(\theta_{ref}) - \chi^2(\theta_{fit})$ values (across all 1000 noise realizations) were below the threshold value ($\alpha = 0.01$). **Any improvement of $\chi^2$ below that of $\Delta_{1-\alpha}$ should be attributed to overfitting rather than to a significant improvement in the hypothesis associated with a model.**

**Implementation of an adaptive stepping algorithm for profile likelihood calculations**

Determining appropriate step sizes is very important when calculating the PPL. When the PPL is steep, small steps should be taken, and when PPL is flat, large steps should be taken. This requires an adaptive stepping procedure. We utilized an approach based on a binary search and by consulting relevant literature[1] to choose appropriate step sizes for each new PPL calulation according to the following steps:

1) Start with the calibrated parameter ($\theta_{i,cal}$), re-optimize all other parameters ($\theta_{j \neq i}$), then calculate ${\chi_{PL}}^2$.

2) Take a step away from the calibrated parameter in the positive direction (starting with the maximum step size), fix the parameter at the new value, re-optimize all other parameters, and calculate the minimum possible $\chi^2$.
   - The step is accepted only if:

$$\chi^2(\theta_{last} + \theta_{step}) - \chi^2(\theta_{last}) \approx q \cdot \Delta_{1-\alpha} \qquad \textbf{(Equation S2)}$$

     where $q = 0.1$, $\theta_{last}$ is the parameter set associated with the previous step, and $\theta_{step}$ is the parameter set associated with the current step.
   - In our implementation, steps within $0.1 \cdot q \cdot \Delta_{\alpha}$ and $2 \cdot q \cdot \Delta_{1-\alpha}$ are accepted.
   - The maximum number of steps in either direction equals 50. Only accepted steps count towards this value.
   - The maximum step size is $0.2 \cdot \theta_{i,cal}$.
   - The minimum step size is $0.01 \cdot \theta_{i,cal}$ for *b, m, e*, and $k_m$. For parameters $k_{bind}$ and *m*, the minimum step size is $0.0001 \cdot \theta_{i,cal}$. Smaller minumum steps are often necessary if the calibrated parameter value is large compared to the minumum parameter bound.

- If the maximum step value is taken and the resulting parameter value is $\leq 0$, then update the maximum step value to be half of the previous value, recalculate the given step value, and then proceed to calculation of the profile likelihood.
    - Repeat this step until the fixed parameter value is > 0. If the minimum step value is reached before the fixed parameter value becomes > 0, then break the current profile likelihood calculation loop and move on to the next direction or fixed parameter.

3) If the step is accepted, try another step using the same step size as the previous step.

4) If the step is not accepted, try another step.
    - If the step was too large (that is, $\chi^2(\theta_{last} + \theta_{step}) - \chi^2(\theta_{last})$ is too big), then try a smaller step.
    - If the step was too small (that is, $\chi^2(\theta_{last} + \theta_{step}) - \chi^2(\theta_{last})$ is too small), then try a larger step.
    - The size of the next step is calculated using a binary search. In the first step, the maximum step is used. If this step is too big, then a step that is halfway in between 0 and the maximum step is tried; for this step, the minimum step bound is set to 0 and the maximum step bound is set to the maximum step such that

$$step_{new} = \frac{min_{bound} + max_{bound}}{2} \quad \text{(Equation S3)}$$

    - Next, the new step value is used to determine the next value of $\theta_i$ to try. If the step is again too large, then the maximum bound is set to the current step value and **Equation S3** is used to propose a new step value. If the step value is too small, then the minimum bound is replaced with this step value and a new step value is proposed again according to **Equation S3**.

5) Keep taking steps until:
    - The parameter bound is reached OR
    - The profile likelihood exceeds $1.1 \cdot \Delta_{1-\alpha}$ OR
    - The maximum number of steps is reached

6) Repeat steps 1–5 in the negative direction.

7) Repeat steps 1–6 for each of the free parameters.

**Avoidance of local minima when calculating the profile likelihood**

The presence of local minima poses a practical challenge for appropriate execution of the PPL approach[3, 4] (**Figure S4**). Each data point along the PPL of each parameter is an individual parameter estimation problem (each with the same model structure, but different free and fixed parameters) that must be solved independently by executing the parameter estimation method. If the parameter estimation method is unable to identify a global minimum for each individual parameter estimation problem, then the PPL will not be interpretable. It is often necessary to tune hyperparameters again when executing the PPL approach (separate from the hyperparameters used in Module 1 and Module 2). In Module 3, hyperparameter tuning can be practically executed by trying different hyperparameter combinations, visualizing the profile likelihood results, and then trying again if necessary (rather than based on some fitting criterion). Hyperparameters should be tuned until an increase in the hyperparameters no longer causes a change in the PPL results, indicating that the PEM has identified a global minimum for each re-optimization. In the case study, Modules 1 and 2 were run with hyperparameters 100 and 10 for the number of parameter sets in the global search and number of initial guesses for optimization, respectively. PPL simulations in Module 3 were run with hyperparameters 1000 and 100. All PPL simulations begin with rerunning Module 2 to identify calibrated parameters using the same hyperparameters as Module 3.

**Supplementary Note 2: Description of functions included in source code and suggestions for adapting the code to new models**

** If the user changes the model, this function must be updated.*
*^ If the user changes the training data, this function must be updated.*
*# If the user changes anything, this function must be updated.*
*Other functions may need to be updated depending on other modeling choices, such as normalization strategy, cost function, or parameter estimation method.*

**`Test.py`** (executable)

- ^ `saveRefData(data)` saves the reference training data as a DataFrame
- ^ `generateRefData(p_ref)` generates a set of reference training data given the parameters `p_ref` and outputs a plot showing the reference model trajectory (without error) on the same plot as the reference training data (with technical error and biological error)
- ^ `testSingleSet(p)` simulates the training data for a given parameter set, `p`, and outputs a plot showing the simulated and experimental (or "true") training data on the same plot

**`Settings.py`**

- # `init()` defines the settings for the run. The variables defined here are imported into other files.
  - Make sure to change the name of the results folder with each new run.

**`DefineExpData.py`**

- ^ `defineExp(args)` defines the experimental data (training data) given a set of `data` and a `model`. Note that the data is structured in a specific way (`x, data, error`) because these variables are directly fed into `LMFit` to perform optimization. Therefore, changing the structure of the data in this file may cause problems with downstream functions.
  - Note that the code refers to training data as experimental data.

**`Solvers.py`**

- `calcRsq(dataX, dataY)` calculates and outputs the $R^2$ between two given datasets, `dataX` and `dataY`.
- `calcChi2(exp, sim, std)` calculates and outputs the weighted `chi2` between two given datasets, `exp` and `sim`, given a list of error values (`std`) for each experimental datapoint. Each of the three inputs must be the same length.
- * `model_AB(y, t, v)` and `model_CD(y, t, v)` define the ODEs to be solved for each datapoint, depending on the model (models A/B and models C/D have slightly different structure due to model reduction). These functions are structured such that they can be used directly with `odeint` (in `solveSingle`) and `LMFit` (in `Run.py`)
- * `solveSingle(args)` solves the model for a single datapoint (single set of component doses) and returns either the simulated reporter expression value at

the final timepoint or the entire solution of all states at all timepoints (if `output = 'save internal states'`)

**`Analysis.py`**

- `analyzeSingleRun(args)` takes in a set of optimization results and plots the CF trajectory (CF vs function evaluation) for each initial guess
- `plotPEMEvaluation(args)` evaluates and plots the PEM evaluation criterion

**`ModelSelection.py`**

- ^ `plotModelSelection(args)` plots the model selection panel for a given set of results folders
  - *If the training data structure changes, then this function will need to be updated.*
  - *The results folder locations for the competing models must be manually input.*

**`Run.py`** (executable)

General parameter estimation/solver code

- ^ * `solveAll(p)` solves for reporter expression at the final timepoint for each datapoint in the dataset, given a set of parameters, `p`
- `solvePar(row)` simulates the entire dataset given a set of parameters, as defined in the input `row`. This function is set up to be directly callable by upstream multiprocessing code to facilitate parallelization of simulations.
- `optPar(row)` enables parameter optimization given the conditions defined in the input `row`. This function is set up to be directly callable by upstream multiprocessing code to facilitate parallelization of simulations.
- ^ `plotTrainingDataFits(df)` takes in a `df` with a set of optimization results for a number of initial guesses and plots the fit to training data

Module 1: code to generate and simulate PEM evaluation data

- ^ `savePemEvalData(args)` saves the generated PEM evaluation data in a specific structure to be used in downstream code
  - This function will need to be updated if the structure of the training data is changed (add an `elif` statement)
- ^ `generatePemEvalData(args)` generates a number of PEM evaluation datasets as defined by `num_pem_eval_datasets` and adds technical error to each dataset
  - This function will need to be updated if the structure of the training data is changed (add an `elif` statement)
- ^ `addNoise(args)` adds technical error individually to each datapoint randomly depending on some predefined distribution of error

- The final section of this function renormalizes the data following the addition of technical error and might needed to be updated if the structure of the training data is changed.
- This function is also used to add technical error to the true data (In `Test.py`) and to the noise realizations (in module 4 in `Run.py`)

- `unpackPemEvalData(args)` unpacks the output from the global search (based on the structure of the output data from upstream multiprocessing code)
- `runGlobalSearchPemEval()` runs the global search only. This function is separate from the optimization function for PEM evaluation data because the same global search can be used for all PEM evaluation datasets.
- `runOptPemEval (args)` runs optimization for a given set of PEM evaluation datasets.
- ^ `defineExpPemEvalData(args)` defines the experimental data for each set PEM evaluation data

Module 2: code for parameter estimation with training data

- `plotParamDistributions(df)` plots the distribution of optimized parameter sets for which $R^2 \geq 0.99$.
- `unpackOutputGS(args)` unpacks the output from the global search (based on the structure of the output data from upstream multiprocessing code)
- `runParameterEstimation(args)` completes the parameter sweep, global search, filtering, and optimization required for a full run of the parameter estimation method

Module 3: code for parameter identifiability analysis

- `calcPL(args)` calculates the parameter profile likelihood for each free parameter
- `plotPL(args)` plots the parameter profile likelihood for each free parameter
- * `plotPL_Consequences(args)` plots the parameter relationships and internal model states along the parameter profile likelihood for each free parameter
- ^ `calcThresholdPL(args)` calculates the threshold PL value and plots the $\chi^2$ distribution

The rest of the code in `Run.py` is used to run each individual module (1, 2, 3) if the module is defined in `Settings.py`. To run the code for each different model, the model variable in `Settings.py` should be set to a string defining the appropriate model, e.g., `'model A'`. The list `real_param_labels_free` should appropriately define the free parameters in the simulation, e.g., `['e', 'm', 'km', 'n']` for model D. The list `p_ref` should also be updated to properly define the reference parameter values for the given model, e.g., for models C and D, *m** takes the place of *m*, and the value of *m** (rather than *m*) should be included in `p_ref`. The code on GitHub contains the `Settings.py` file for model D, which is the only model with all identifiable parameters and has the fastest run time (unidentifiable parameters significantly increase run time).

**Supplementary Note 3: Description of simulation outputs and source data**
Note: Capitalized headings refer to folder names in the source data.

**GAMES**

- REFERENCE TRAINING DATA.xlsx includes the reference training data generated with reference parameters
- REFERENCE TRAINING DATA.svg shows the reference training data generated with reference parameters

**TEST**

- FIT TO TRAINING DATA.svg shows the simulation results for a given parameter set, p, on the same plot as the training data

**MODULE 1 - PARAMETER ESTIMATION WITH PEM EVALUATION DATA**

**GENERATE PEM EVALUATION DATA**

- CONDITIONS.txt includes the conditions used to start the run
- PARAM SWEEP.xlsx includes the parameters (parameters only and not simulation results) to be used in the global search, as generated via Latin Hypercube Sampling.
- GLOBAL SEARCH RESULTS. xlsx includes the global search results (based on agreement with experimental data)
- INITIAL GUESSES.xlsx includes the PEM evaluation parameter sets (filtered global search results based on agreement with experimental data)
- PEM EVALUATION DATA RAW.xlsx includes the PEM evaluation training datasets (before noise is added)
- PEM EVALUATION DATA NOISE.xlsx includes the PEM evaluation training datasets (after noise is added)

**PARAMETER ESTIMATION WITH PEM EVALUATION DATA**

- PARAM SWEEP.xlsx includes the parameters (parameters only and not simulation results) to be used in the global search, as generated via Latin Hypercube Sampling.
- GLOBAL SEARCH RESULTS.xlsx includes the global search results (based on agreement with experimental data). We note that the same global search is used for all PEM evaluation datasets. The cost function is calculated individually for each PEM evaluation dataset and initial guesses for optimization are chosen for each individual PEM evaluation dataset accordingly.
- RUN x/
    - Folder including parameter estimation results for PEM evaluation dataset, x.
    - CONDITIONS.txt holds the conditions used to start the run
    - INITIAL GUESSES.xlsx includes the filtered data (after the global search) to be used as initial guesses.

    - OPT RESULTS.xlsx includes the optimization results (the final step of parameter estimation method)
    - FITS.svg plots the PEM evaluation dataset and simulations with optimized parameter values (across multiple initial guesses)
    - BEST FIT.svg plots the PEM evaluation dataset with the best case fit from optimization (parameter set yielding the lowest cost function, $\chi^2$)
- ANALYSIS/
    - CF TRAJECTORY PLOTS RUN x.svg plots the $\chi^2$ trajectories for each initial guess along each function evaluation and shows the initial and optimized (final) $\chi^2$ values for each initial guess. This plot can be very useful for debugging the PEM.
    - PEM EVALUATION CRITERION.svg plots the PEM evaluation criterion across all PEM evaluation datasets
        - PEM EVALUATION CRITERION R2 >= 0.90.svg plots the PEM evaluation criterion across all PEM evaluation datasets and scales the y-axis from 0.90 to 1.0.

**<u>MODULE 2 – FIT TO EXPERIMENTAL DATA</u>**

- CONDITIONS.txt includes the conditions used to start the run, as defined in Settings.py.
- PARAM SWEEP.xlsx includes the parameters (parameters only and not simulation results) to be used in the global search, as generated via Latin Hypercube Sampling.
- GLOBAL SEARCH RESULTS.xlsx includes the global search results before filtering.
- INITIAL GUESSES.xlsx includes the filtered data (after the global search) to be used as initial guesses.
- OPT RESULTS.xlsx includes the optimization results (the final step of the parameter estimation method).
- FITS.svg plots the training data and simulations with optimized parameter values (across multiple initial guesses).
- BEST FIT.svg is a plot comparing the training data with the best case fit from optimization (parameter set yielding the lowest cost function, $\chi^2$).

**<u>MODULE 3 – PARAMETER IDENTIFIABILITY ANALYSIS</u>**

- CHI2 DISTRIBUTION.svg shows the results of a simulation study to verify the $\chi^2$ distribution for the given model structure and free parameters and shows the practically determined threshold value
- CONDITIONS PL.txt includes the conditions used for the PPL calculations, including the threshold value and the starting, calibrated parameter set.
- PROFILE LIKELIHOOD PLOTS.svg plots the profile likelihood of each free parameter in subplot form.
- PROFILE LIKELIHOOD RESULTS x.xlsx includes the data used to generate the plots in PROFILE LIKELIHOOD PLOTS.svg.
    - Analogous files are included for each free parameter (represented here as x).

- INTERNAL STATES ALONG x.svg plots the trajectories of the internal model states (for a single dose combination of 50 ng activation domain, 50 ng DNA-binding domain and a saturating dose of ligand of 100 nM) along the free parameter, x.
    - Analogous files are included for each free parameter (represented here as x).
- PARAMETER RELATIONSHIPS ALONG x.svg plots the relationships between the free parameter, x, and all other parameters (along the profile likelihood)
    - Analogous files are included for each free parameter (represented here as x).
- INITIAL GUESSES.xlsx, and PARAM SWEEP.xlsx and OPT RESULTS.xlsx show intermediate results of the most recent PEM run in the profile likelihood calculations. These files can be ignored in module 3.

### Supplementary Figures

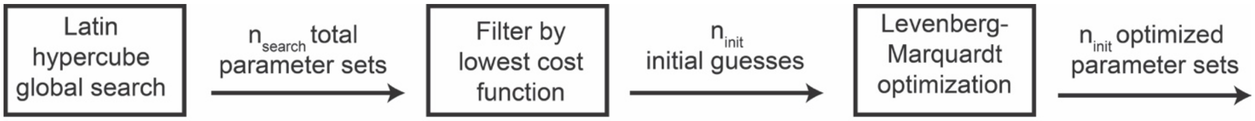


**Figure S1. Description of the parameter estimation method used throughout this tutorial.** A global search is used to identify parameter sets to use as initial guesses for an optimization algorithm.

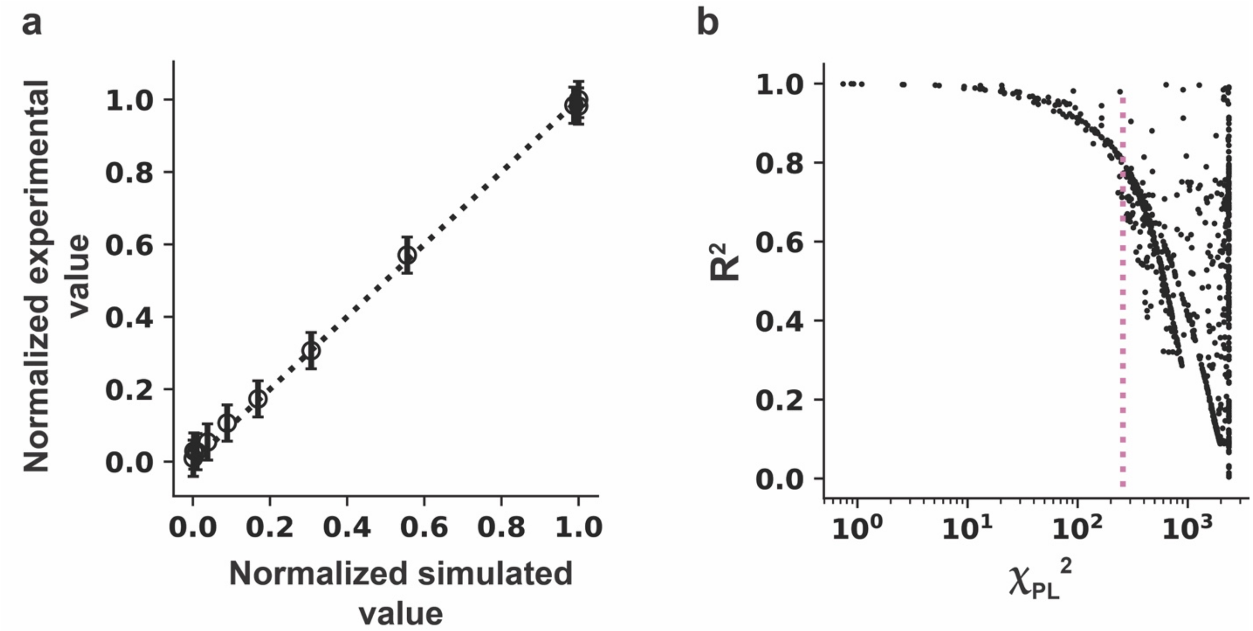


**Figure S2. Rationale for using $R^2$ to quantify goodness of fit. (a)** Representative example of a parity plot depicting the correlation between the experimental and simulated values. This correlation is assumed to be linear, and for good fits the points will ideally fall along the line y = x (black dotted line). **(b)** Representative example of the correlation of $R^2$ and $\chi^2$ using a random 1000 parameter set global search. $R^2$ and $\chi^2$ are closely negatively correlated for parameter sets with $\chi^2$ values in the bottom 10$^{th}$ percentile (to the left of the pink dotted line).

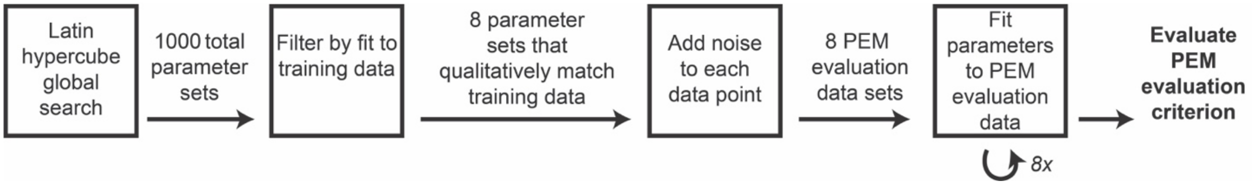


**Figure S3. Description of the generation of simulated data and subsequent determination of the PEM evaluation criterion.** A global search is used to identify 8 parameter sets that qualitatively match the training data. Technical and biological measurement error are added to each data set to generate 8 PEM evaluation data sets, as described in the main text. Each of the 8 new data sets are independently used to define 8 parameter estimation problems. After fitting parameters to each PEM evaluation data set, the PEM evaluation criterion can be determined.

| Failure mode | Example | Illustration | Suggested remediation |
|---|---|---|---|
| The PPL for a parameter $\theta_i$ is **not evaluated over the entire range of plausible parameter values** based on the parameter bounds. | If the calibrated value of $\theta_i$ is small relative to the value at which the PPL reaches the threshold, then the maximum step size will be relatively small and the maximum number of steps may be reached before the PPL reaches the threshold. | 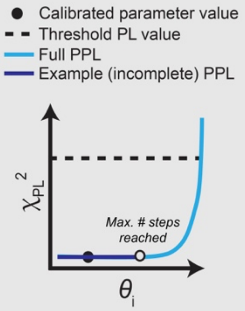 | Tune PPL hyperparameters, such as the maximum number of steps and the maximum step size until the entire range of values is evaluated. This should be done carefully and individually for each parameter, as there is a tradeoff between speed and properly evaluating every region of the PPL plot. |
| The PPL for a parameter $\theta_i$ is **bumpy with no well-defined shape.** | If the PEM is identifying local minima at some $\theta_i$ values, rather than the global minimum, then the resulting PPL may appear bumpy with jumps in $\chi_{PL}^2$, rather than the expected smooth curve. | 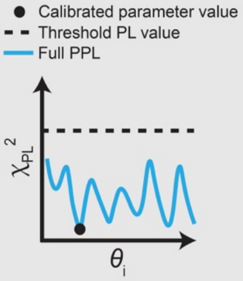 | Increase PEM hyperparameters, such as the number of parameter sets in the global search and the number of initial guesses for optimization, until PPLs appear smooth. The PPL often requires hyperparameters that are more conservative (higher values, in this case) than required for estimating parameters. |
| The PPL for a parameter $\theta_i$ has **bounds that are unknown, infinite, or span a very large range.** *This is distinct from the first failure mode, in which the PPL is given parameter bounds but is unable to evaluate the entire given range.* | If parameter bounds are infinite, it will not be possible to evaluate the PPL across this entire range. Evaluating over a large range may also be difficult and time-consuming. For example, if the calibrated value is small and the upper parameter bound is many orders of magnitude larger, PPL calculations at high $\theta_i$ values may take a long time. This can sometimes be mitigated by tuning PPL hyperparameters. | 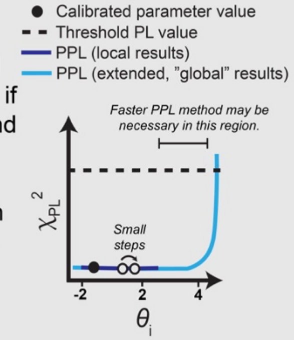 | Use an a priori method to refine structurally unidenfitiable parameters, then use the PPL to investigate other parameters[3, 12]. Set reasonably large parameter bounds (-3 to 3 in orders of magntiude from the calibrated parameter, for example) for parameters with infinite bounds. For models with a large number of unidentifiable parameters, a faster PPL method may be required[3, 11, 12]. |
| The PPL for a parameter $\theta_i$ **is impacted by the bounds of another parameter** $\theta_j$. | Consider two parameters $\theta_i$ and $\theta_j$ that compensate for one another. If the fixed value of $\theta_i$ is increased and the value of $\theta_j$ has already reached its upper bound, then $\theta_j$ will no longer be able to compensate for changes in $\theta_i$. Therefore, $\theta_i$ may appear to be practically unidentifiable, although the parameter would be expected to be classified as structurally unidenfitiable if parameter bounds were not taken into account, for example in a global analysis. | 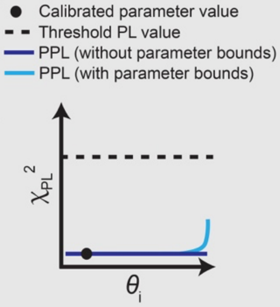 | Plots of parameter relationships associated with each PPL can be used to interpret results and detemine next steps. An example of this is shown in the results for Model B in Module 3. If results cannot be justified (such as if many parameters compensate for other another and these relationships are not clear), the modeler can use an a priori method[3, 12] to refine structurally unidenfitiable parameters, then use the PPL to investigate other parameters. |

**Figure S4. Module 3 failure modes, illustrations, and suggested remediation strategies.** In practice, any number of these strategies may be necessary to successfully evaluate the PPL for a given parameter in a given model. Additional failure models and suggested remediation strategies are included in Supplementary Table 3.

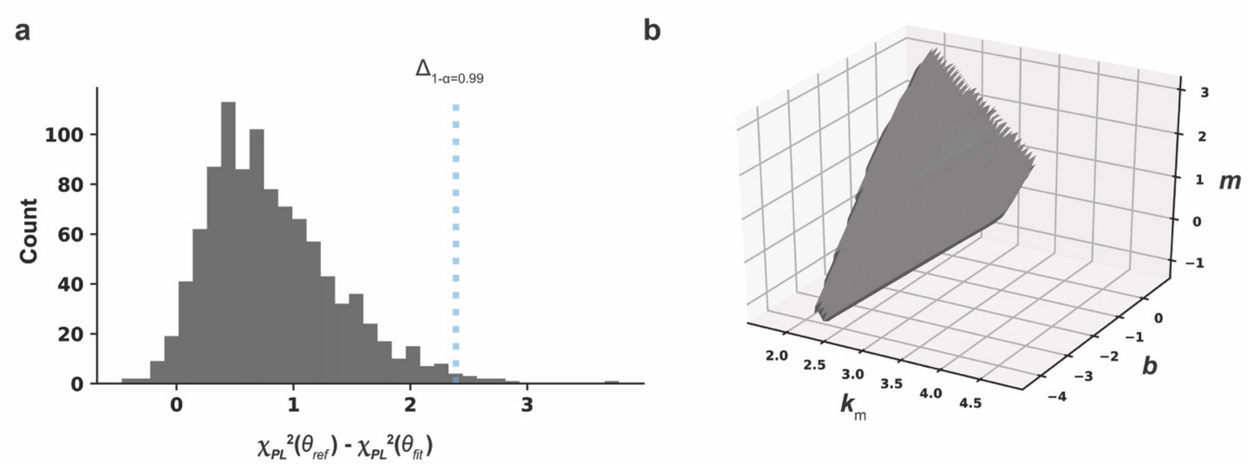


**Figure S5. Additional PPL results for Model A. (a)** Evaluation of the $\chi^2$ distribution via a simulation study. 1000 individual noise realizations were generated. Parameters were individually estimated for each of the noise realizations to calculate $\chi^2(\theta_{fit})$. Reference parameters were used to calculate $\chi^2(\theta_{ref})$. The difference between these values, $\chi^2(\theta_{ref}) - \chi^2(\theta_{fit})$, represents the amount of overfitting for each noise realization. $\Delta_{1-\alpha}$ (blue dotted line) was determined by evaluating the 99% confidence interval ($\propto$ = 0.01) of the distribution ($\Delta_{1-\alpha} = 2.4$). **(b)** Three-dimensional plot of $k_m$, $b$, and $m$ along the unidentifiability associated with $m$. The surface is smooth, indicating dependencies between the three parameters. The logarithm ($\log_{10}(\theta)$) of each parameter is plotted.

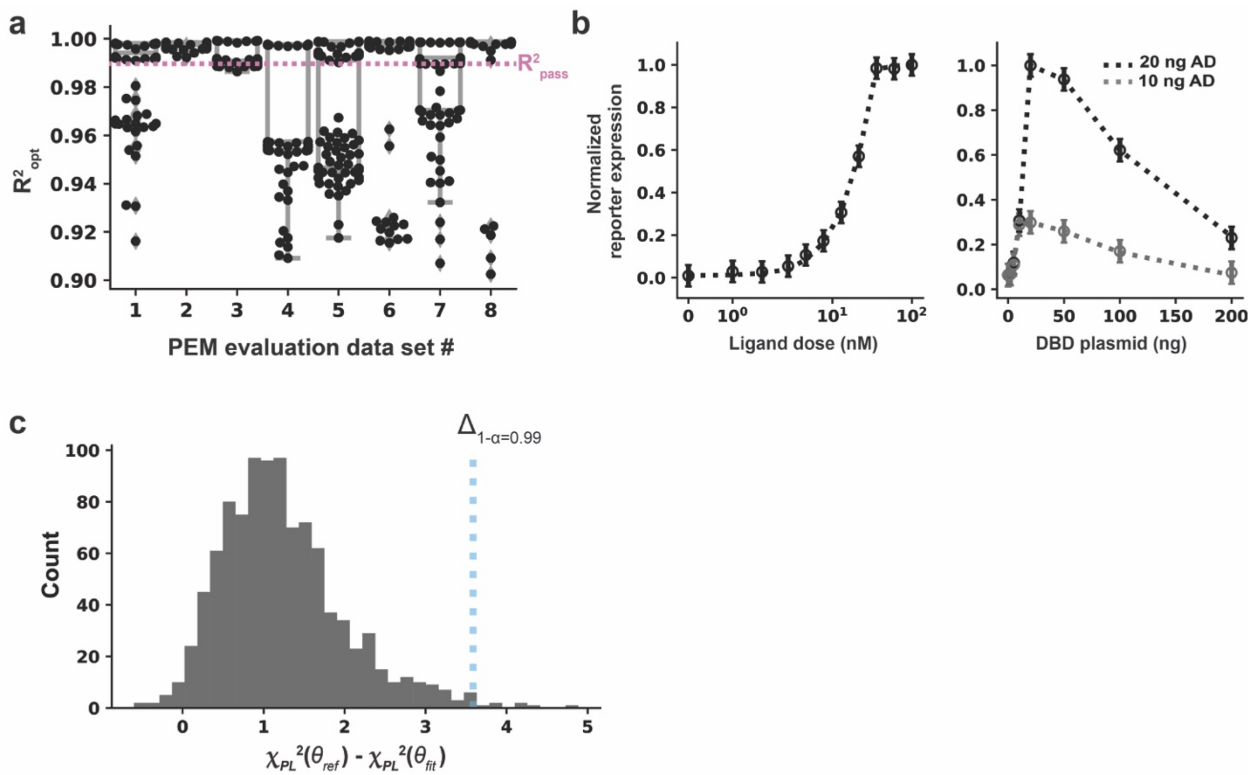


**Figure S6. PEM evaluation, parameter estimation, and determination of confidence threshold for Model B. (a)** PEM evaluation criterion with 1000 parameter sets in the global search and 100 initial guesses. Results are shown only for parameter sets yielding $R^2 \geq 0.90$. The PEM evaluation criterion is satisfied. **(b)** Best fit to the training data using the calibrated parameter set**.** The visual inspection criterion is satisfied. Parameter values are in Supplementary Table 2. **(c)** Determination of the confidence threshold for PPL calculations ($\Delta_{1-\alpha}$ = 3.6).

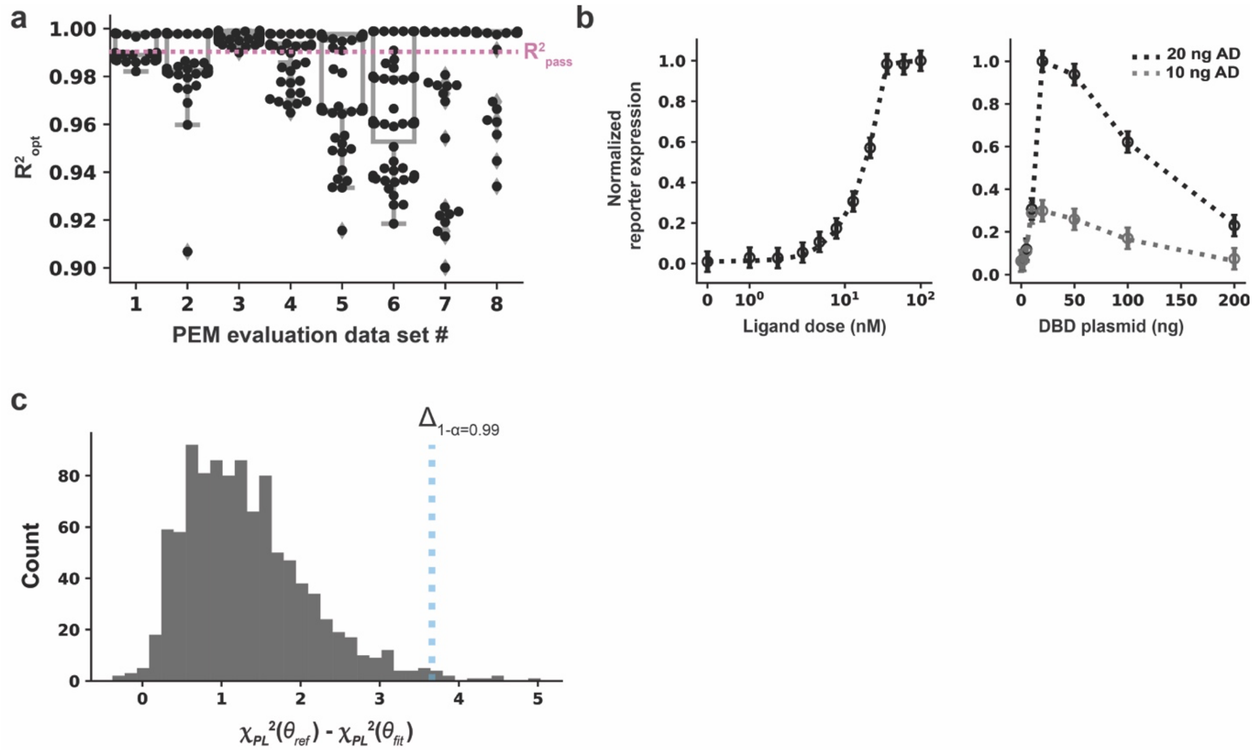


**Figure S7. PEM evaluation, parameter estimation, and determination of confidence threshold for Model C. (a)** PEM evaluation criterion with 1000 parameter sets in the global search and 100 initial guesses. The PEM evaluation criterion is satisfied. **(b)** Best fit to the training data using the calibrated parameter set**.** The visual inspection criterion is satisfied. Parameter values are in Supplementary Table 2. **(c)** Determination of the confidence threshold for PPL calculations ($\Delta_{1-\alpha}$ = 3.7).

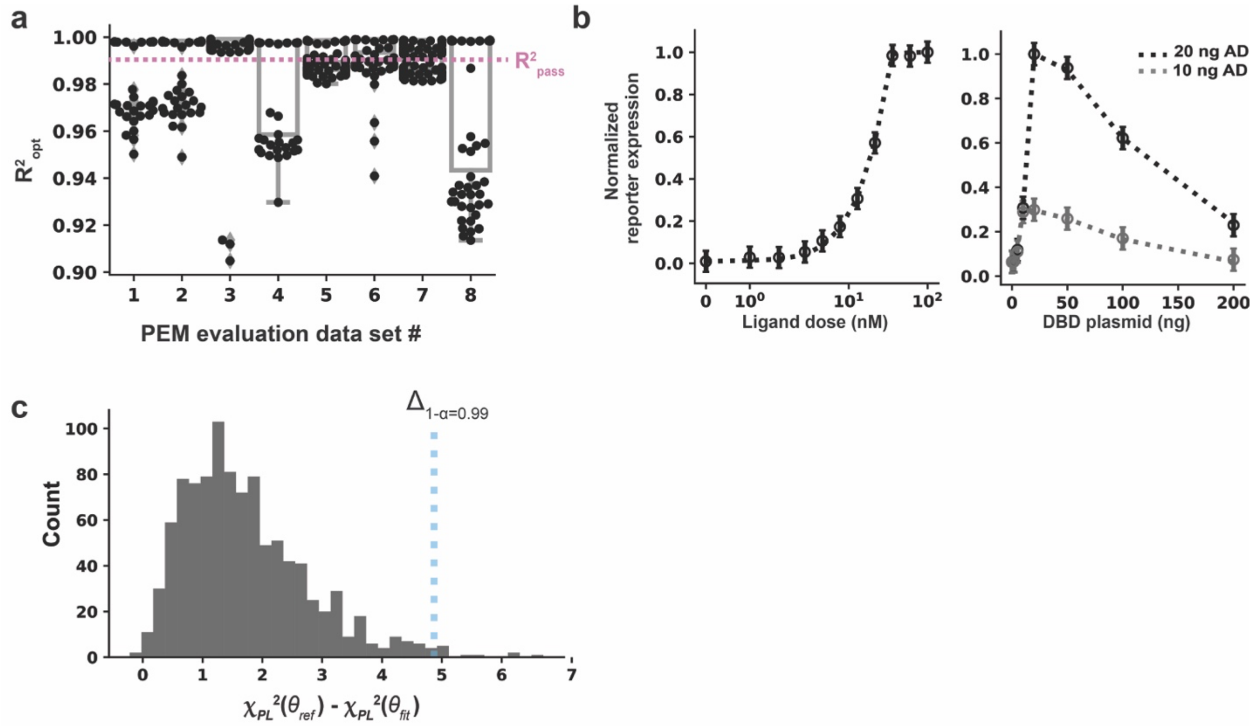


**Figure S8. PEM evaluation, parameter estimation, and determination of confidence threshold for Model D. (a)** PEM evaluation criterion with 1000 parameter sets in the global search and 100 initial guesses. The PEM evaluation criterion is satisfied. **(b)** Best fit to the training data using the calibrated parameter set**.** The visual inspection criterion is satisfied. Parameter values are in Supplementary Table 2. **(c)** Determination of the confidence threshold for PPL calculations ($\Delta_{1-\alpha}$ = 4.9).

### Supplementary Tables

#### Supplementary Table 1. Kinetic parameters for the case study ODE model^

| Parameter | Free/fixed | Value | Description | Reference |
|---|---|---|---|---|
| $e$ | Free | n.a. | Conversion factor to compare ligand dose in experimental units to ligand dose in model units | n.a. |
| $b$ | Free | n.a. | Background promoter activation | n.a. |
| $k_{\mathrm{bind}}$ | Free | n.a. | Binding constant of DNA-binding domain, activation domain, and ligand | n.a. |
| $m$ | Free | n.a. | Maximal expression from the promoter | n.a. |
| $k_{\mathrm{m}}$ | Free | n.a. | Promoter activation coefficient | n.a. |
| $n$ | Free | n.a. | Hill coefficient (degree of cooperativity) | n.a. |
| $k_{\mathrm{txn}}$ | Fixed | 1 U | Transcription rate constant for all mRNA species | [5] |
| $k_{\mathrm{trans}}$ | Fixed | 1 U | Translation rate constant for all protein species | [5] |
| $k_{\mathrm{deg,mRNA}}$ | Fixed | 2.7 $\mathrm{hr}^{-1}$ | Degradation rate constant for all mRNA species | [5] |
| $k_{\mathrm{deg,protein}}$ | Fixed | 0.35 $\mathrm{hr}^{-1}$ | Degradation rate constant for all protein species except the reporter | [5] |
| $k_{\mathrm{deg,reporter}}$ | Fixed | 0.029 $\mathrm{hr}^{-1}$ | Degradation rate constant for all protein species except the reporter | [5] |
| $k_{\mathrm{deg,ligand}}$ | Fixed | 0.01 $\mathrm{hr}^{-1}$ | Degradation rate constant for the ligand | Arbitrarily low |

^U refers to an arbitrary transcriptional unit as described previously[5, 6]. n.a. means not applicable for free parameters.

**Supplementary Table 2. Kinetic parameter values for the case study ODE model^**

| Parameter | | | Calibrated value | | | | |
|---|---|---|---|---|---|---|---|
| | Units | Reference value | Model A 100 GS + 10 OPT | Model A 1000 GS + 100 OPT | Model B 1000 GS + 100 OPT | Model C 1000 GS + 100 OPT | Model D 1000 GS + 100 OPT |
| $\boldsymbol{e}$ | U nM$^{-1}$ | 15 | 14.5 | 14.6 | 15.3 | 15.3 | 15.2 |
| $\boldsymbol{b}$ | n.a. | 0.05 | 0.001 | 0.008 | 29.3 | 1 | 1 |
| $\boldsymbol{k}_{\mathbf{bind}}$ | U$^{-2}$ t$^{-1}$ | 0.05 | 0.462 | 0.235 | 0.126 | 0.128 | 1 |
| $\boldsymbol{m}$ | n.a. | 36 | 3.081 | 146.0 | 17071 | | |
| $\boldsymbol{k}_{\mathbf{m}}$ | U | 100 | 355 | 1792 | 103 | 103 | 105 |
| $\boldsymbol{n}$ | n.a. | 2 | 1.41 | 1.50 | 2.03 | 2.03 | 2.04 |
| $\boldsymbol{m}^{*}$ | n.a. | 720 | | | | 582 | 586 |

^U refers to an arbitrary transcriptional unit as described previously[5, 6]. Gray shading indicates that a parameter is not included in the given model (no value in cell) or the fixed parameter data (value in cell).

### Supplementary Table 3. Module failure modes and suggested remediation

| Module 1: Evaluate parameter estimation method | |
|---|---|
| **Observation** | **Suggested remediation** |
| The PEM evaluation criterion is not met. | 1) Look for implementation errors in the code. This can be investigated by trying a much simpler, toy parameter estimation problem, such as estimating a single parameter, and by analyzing the cost function trajectory (cost function vs. function evaluation) for the optimization step of the PEM (if implementation is correct, CF should decrease with # of function evaluations).<br>2) Investigate different normalization strategies[7].<br>3) Tune hyperparameters[8].<br>4) Try a different PEM by consulting the literature and proposing a method that is well-suited to the cost function landscape[7 9]. For example, if the data are noisy, a PEM using a dynamic recursive estimator might be appropriate[10].<br>5) If an increase in hyperparameters does not have an effect on the PEM results and changing the PEM to a different method does not have an effect on the PEM results, then the PEM evaluation criterion may need to be lowered. |
| The measurement error in the training data is not approximately normally distributed (for example, if data from two different experiments vary in order of magnitude). | Alter the cost function to use a different error metric that is approximately normally distributed, such as the root mean squared error (RMSE). |
| The PEM cannot be executed because the ODE solver is unable to solve the equations for some or all parameter sets (e.g., the code gets stuck when trying to solve the ODEs) or takes a prohibitively long time to solve the equations. | Try a different ODE solver. For example, if the trajectories are stiff, meaning there are some very slow and very fast processes in the system, Python's solve_ivp with the backward differentiation formula (BDF) algorithm should be used. |

| Module 2: Fit parameters with training data | |
|---|---|
| **Observation** | **Suggested remediation** |
| Dynamic trajectories of state variables are non-physical (e.g., negative values) | 1) Check conservation of mass in the system to make sure equations are written correctly.<br>2) Rescale parameters or initial conditions if orders of magnitude vary greatly across the set of parameters or the set of initial conditions.<br>3) Try a different ODE solver (e.g., a solver that is designed to handle stiff equations, if applicable) |

| | |
|---|---|
| The model does not describe one or more features of the data. | 1) Return to module 0 and propose a new model with additional or different mechanisms.<br>2) Consider different normalization strategies. |
| **Module 3: Assess parameter identifiability** | |
| **Observation** | **Suggested remediation** |
| Profile likelihoods are bumpy and uninterpretable. | Increase PEM hyperparameters. |
| Profile likelihoods are not evaluated appropriately for some parameter ranges (e.g., steps in the negative direction are too large and therefore multiple orders of magnitude of the fixed parameter value are skipped over in successive steps or steps in either direction are too small such that the entire range of the fixed parameter value, from minimum to maximum parameter bound is not evaluated). | Tune PPL hyperparameters: Minimum and maximum step sizes, total number of steps in either direction, the aspired ${\chi_{PL}}^2$ increase of each step ($q$) (see **Supplementary Note 1**). |
| One or more parameters are unidentifiable and do affect model predictions. | Return to module 0 and collect additional experimental data, such as the concentration of intermediate states or higher resolution sampling, if the timescales of the unidentified terms are not resolved. |
| One or more parameters are unidentifiable and do not affect model predictions. | Return to module 0 and propose a model reduction strategy, based on which parameters compensate for one another. |
| Profile likelihood calculations are computationally infeasible with the algorithm used here (e.g., for large models or those with many structurally unidentifiable parameters). | 1) Parallelize calculations using a supercomputer cluster[4].<br>2) Use an a priori method for assessing structural unidentifiability[3, 11, 12].<br>3) Use a progressive step choice algorithm, based on the first and second derivatives of the cost function, for profile likelihood calculations[1, 13].<br>4) Try a simpler model or increase the training data, as the computational expense may be due to most parameters being unidentifiable. |
| **Module 4: Compare candidate models** | |
| **Observation** | **Suggested remediation** |
| The selected model does not adequately predict the test data. | Return to module 0 and propose a new model which includes more mechanisms or different mechanisms based on the observations. Consider including the test data as training data and collect a new set of test data. |
